# Monomeric amyloid *β*-peptide (1-42) significantly populates compact fibril-like conformations

**DOI:** 10.1101/2020.06.23.156620

**Authors:** Bogdan Barz, Alexander K. Buell, Soumav Nath

## Abstract

The aggregation of the amyloid *β* (A*β*) peptide is a major hallmark of Alzheimer’s disease. This peptide can aggregate into oligomers, proto-fibrils, and mature fibrils, which eventually assemble into amyloid plaques. The peptide monomers are the smallest assembly units, and play an important role in most of the individual processes involved in amyloid fibril formation, such as primary and secondary nucleation and elongation. The structure of the A*β* monomer has been shown to be very dynamic and mostly disordered, both in experimental and in computational studies, similar to a random coil. This structural state of the monomer contrasts with the very stable and well defined structural core of the amyloid fibrils. An important question is whether the monomer can adopt transient fibril-like conformations in solution and what role such conformations might play in the aggregation process. Here we use enhanced and extensive molecular dynamics simulations to study the A*β*42 monomer structural flexibility with different force fields, water models and salt concentrations. We show that the monomer behaves as a random coil under different simulation conditions. Importantly, we find a conformation with the N-terminal region structured very similarly to that of recent experimentally determined fibril models. This is to the best of our knowledge the first monomeric structural ensemble to show such a similarity with the fibril structure.

## Introduction

The molecular origins of Alzheimer’s Disease (AD) are linked to the accumulation of abnormally folded proteins and peptides such as tau and amyloid *β* (A*β*) into intracellular neuronal tangles and extracellular amyloid plaques, respectively. The two main A*β* variants found in the brain differ only at the C-terminal two amino acids, with A*β*40 being more abundant than A*β*42 (~5-10%). In AD patients, however, the extracellular concentration of A*β*42 has been shown to be considerably increased. ^1,2^ The formation of amyloid fibrils is a complex process, including primary and secondary nucleation reactions, as well as fibril elongation^3^ and A*β* monomers take part in most of these individual steps. While in typical *in vitro* experiments the monomer is depleted during the aggregation reaction, it is constantly being produced *in vivo* and therefore all monomer-dependent steps are likely to be important during the entire duration of the disease. It is thus very important to characterize the monomer structure in order to understand how its sequence encodes the tendency to form larger assemblies. The A*β* peptide originates from the amyloid precursor protein (APP),^4^ a large glycoprotein which consists of extracellular, trans-membrane and cytoplasmic domains. Upon cleavage by the *γ* secretase protein,^5^ the A*β* peptide is formed with its N-terminus originating from the extracellular domain of the APP, and the C-terminus from the transmembrane domain.^6^ The A*β*42 monomer freshly cleaved from the APP has a considerable amount of helical structure. 3D structures published thus far indicate that A*β*42 adopts a mostly helical conformation in apolar micro-environment^7^ which starts to vanish as soon as polar solvent is added. ^8^ A very important question is related to the monomer structure in polar solvents and how it influences the A*β* aggregation process. Among recent experiments of A*β*42 monomers in phosphate buffer solution are nuclear magnetic resonance (NMR) measurements of various parameters, such as chemical shifts and J-couplings. ^9^ This study reveals a random coil type of conformational ensemble without high propensities for *β*-strand or helical structures for both A*β*42 and A*β*40. The random coil behaviour was also confirmed by circular dichroism (CD) spectroscopy experiments where the A*β* monomers adopt between 12–25% *β*-sheet and much lower helical content.^10,11^ In terms of overall dimensions, the hydrodynamic radius of A*β*40 was measured to be ~1.6 nm at room temperature^12^ while for A*β*42 it adopts a range of values between 0.9–1.8 nm.^13–17^ Computationally, the A*β*42 monomer structure has been studied with various simulating techniques and force fields. ^18^ The general picture that emerges from the numerous molecular dynamics (MD) simulation studies on A*β* monomer is that the observed structural characteristics are very diverse and highly dependent on the simulation conditions. ^18^ Recently, modern force fields tailored for intrinsically disordered proteins start producing conformational ensembles more and more similar to those observed in experimental studies. ^19,20^

Here we report structural ensembles of A*β*42 monomer observed with enhanced MD simulations which reproduce experimental NMR observables with increased accuracy compared to previous computational studies as well as expected secondary structure propensities. Other structural characteristics such as the radius of gyration, end-to-end distance and solvent accessible surface area are in agreement with many computational studies, while differing from others. The most remarkable feature of our simulations is the occurrence of a significantly populated set of conformations which has many structural features in common with recent experimental fibril models. We divide this study in two parts. In the first part we show that the structural ensemble sampled in our simulations is overall similar to a random coil, as observed in experimental and other computational studies. In the second part we discuss in detail the fibril-like monomer conformation observed here and how it might relate to the A*β*42 aggregation.

## Results and discussion

Here we present results from three extensive Hamiltonian replica exchange MD (H-REMD) simulations under different conditions: 1) the Amber99SB*-ILDN force field ^21,22^ and the TIP4P-D water model;^23^ 2) the Charmm36m force field^24^ and a modified Charmm TIP3P water model with increased protein-water interaction (labeled Charmm36mW); and 3) the Charmm36mW force field with modified Charmm TIP3P water model and 150 mM NaCl. We also performed a H-REMD simulation with the Charmm36m force field and the original Charmm TIP3P water model which was not included in the detailed analysis and results due to the lack of convergence. Details regarding the force fields, simulation lengths, and the analysis are provided in the Methods section.

### A*β*42 monomers adopt a random coil structure in water

^3^*J_HNHα_* **NMR couplings** NMR couplings are often used for describing the local structure of intrinsically disordered proteins. Recently, Roche *et al.*^9^ measured various NMR parameters of A*β*40 and A*β*42 in solution and concluded that both peptides lack a stable structure and behave very similarly to a random coil. We calculated the ^3^*J_HNHα_* NMR couplings from the three simulations (Amber99SB*-ILDN, Charmm36mW and Charmm36mW with NaCl), as described in the Methods section, and compared these with experimental values.^9^ For a quantitative comparison we use the reduced *χ*^2^ quantity. Overall, the three force fields give reasonable *χ*^2^ values, Amber having the largest deviations from experiment with *χ*^2^ = 3.4, followed by Charmm36mW with *χ*^2^ = 3.1 and Charmm36mW with added salt, which has the best agreement with experiments with a *χ*^2^ = 2.5. The results for the Charmm36mW with 150 mM NaCl are shown in Figure 1 Top, and for the other two cases in Figure S2. The good agreement between the simulated and experimental ^3^*J_HNHα_* couplings for Charmm36mW with NaCl is an improvement compared to reported *χ*^2^ values from other computational studies. ^19,20^ The largest deviations from experimental values occur at amino acids D7, D23 and A42. Four of the Glycines (G9, G25, G29 and G33) have values slightly lower than the experimental ones, while V12, G15, V18, P19 and A30 overestimate the experimental values. The tendencies observed here are closely related to the *β*-sheet propensity of the amino acids, as discussed below. Interestingly, in the case of Amber99SB*-ILDN, the best agreement with experimental J-couplings occurs at the C-terminus for amino acids G25–I41, while in the case of Charmm36mW the agreement is better for residues in the middle region, between E11–A30. For Charmm36mW with NaCl, except for D7 which shows a large deviation, most of the values are close to their experimental equivalents, explaining the low *χ*^2^ value. Of notable importance are amino acids with J-couplings values above 7.5 Hz, corresponding to dihedral angles found in *β*-strand structures. In Figure 1 Top, the central hydrophobic cluster amino acids V18–F20 and the C-terminal V39–I41, have values close or above 7.5 Hz indicating that they are good candidates for forming *β*-sheets. This behavior, also observed in previous experimental studies, ^18,25,26^ is of crucial importance for the fibril-like conformation discussed in the second part of this work. Thus, the *χ*^2^ values confirm that, within simulation and experimental errors, the A*β*42 monomer behaves under three different simulation protocols as a random coil, in agreement with experiments.

**Figure 1:**
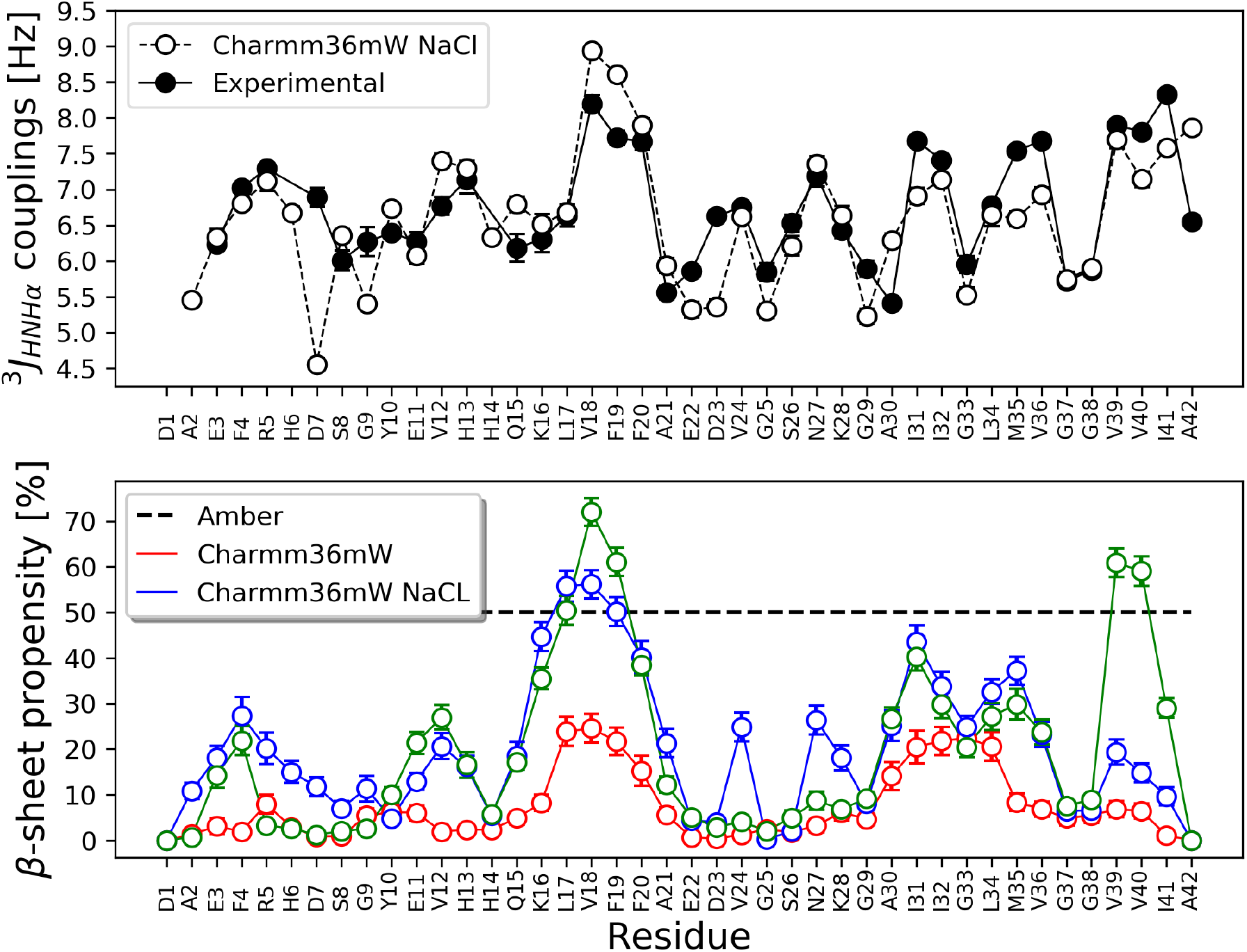
NMR scalar couplings and secondary structure. Top - ^3^*J_HNHα_* NMR scalar couplings calculated for each amino acid for Charmm36mW force field with 150 mM NaCl. Experimental values are shown as black circles and those obtained from the simulations as white circles. Bottom - *β*-sheet propensity per amino acid for the three simulation conditions. The error bars for the simulation data in both plots were calculated using block averaging

#### Secondary structure

The secondary structure propensity per amino acid provides complementary structural information to the J-coupling analysis. As discussed in the methods section, the dominant secondary structure in the trajectories used for analysis is *β*-sheet. We have thus calculated the *β*-sheet propensity per amino acid for each of the three simulation as shown in Figure 1 Bottom. The three cases display a similar main pattern, which is the presence of meta-stable *β*-sheets for the central hydrophobic cluster (the amino acids L17-A21) and the hydrophobic C-terminus (A30-I41), with different propensities. High *β*-strand propensities for L17-A21 and I31-V36 have been observed previously in several computational studies. ^18^ This indicates that overall the sampled conformations have similar structural features between different simulation conditions.

Amber99SB*-ILDN has the smallest propensities with a total average of 7.3±1.1 % *β*-sheet. Both simulations with Charmm36mW force field have higher *β*-sheet propensities with averages of 19.8±2.3 without NaCl and 19.6±2.9 with NaCl. The values from the Charmm36mW simulations are in good agreement with the experimental value of 25.2% reported for A*β*40 monomers.^11^ The largest *β*-sheet propensity of all simulations occurs for Charmm36mW with NaCl at the central hydrophobic cluster, reaching 72% and 61% for V18 and F19, respectively. This is in agreement with the observation that in NMR experiments amino acids V18, F19 and F20 are the residues most likely to have extended conformations. ^9^ The simulation without NaCl has very large propensities at L17-V18, with both residues reaching 56%. The C-terminal region V39-I41 is especially interesting in the case of Charmm36mW with NaCl because it shows very large propensities for V39 (61%) and V40 (59%). Without NaCl, these amino acids have *β*-sheet propensities below 20% for both Charmm36mW and Amber99SB*-ILDN.

Despite the fact that the simulations with Charmm36mW have the highest *β*-sheet propensities, the ^3^*J_HNHα_* couplings are in good agreement with experimental values. Roche *et al.*^9^ showed that A*β*42 monomers lack any long lasting secondary structure, as long as the propensity of these elements is below 50%. With few exceptions, the amino acids in our simulations do have *β*-sheet propensities below 50%, which suggests that the significant amount of *β*-sheet content is subject to constant fluctuations and the conformational ensemble is very diverse. This is also confirmed by the average percentage of *β*-sheet which is in agreement with experimental values that are compatible with a random coil. Additional properties of the conformational ensemble which support the random coil behavior as well as the clustering results are discussed below.

#### Radius of gyration, end to end distance and solvent accessible surface area

In order to further characterize the conformational diversity of the monomeric A*β*, we have calculated distributions for the radius of gyration of the three types of simulations and show them in Figure S3. The distributions have similar shapes, with a strong peak between 1 and 2 nm, and a large shoulder that extends up to 3 nm. These features are more pronounced for the simulations with the Charmm36mW force field. The average radius of gyration values from Table T1 indicate less compact conformations for Amber99SB*-ILDN with an average value of 1.6 nm and more compact conformations for the Charmm36mW force field with or without NaCl with an average value of 1.4 nm in both cases. The values for the Amber99SB*-ILDN are in good agreement with recent studies using similar Amber force fields and water models from Meng *et al.*^19^ and Lincoff *et al.*,^20^ while for Charmm36mW are slightly smaller than the ones from Lincoff *et al.*^20^ Using an empirically parametrized equation which relates the radius of gyration to the hydrodynamic radius for intrinsically disordered proteins,^27^ we have determined hydrodynamic radius distributions for the three types of simulations. The average values from the three distributions are all close to 1.7±0.1 nm, in very good agreement with experimental values of both A*β*40, i.e. 1.6 nm,^12^ and the upper range of values for A*β*42, i.e. 0.9–1.8 nm.^13–17^ The average end to end distance follows a similar trend as the radius of gyration, with a value of 3.2 nm for Amber99SB*-ILDN and 2.7 and 2.8 nm for Charmm36mW without and with NaCl, respectively. The wide distributions of the end to end distance, shown in Figure S3, and the standard deviations of ~1 nm indicate similar types of structural fluctuations in all three simulations. This is also confirmed by the average SASA values of 49 nm^2^ for Amber99SB*-ILDN and 45 nm^2^ (44 nm^2^) for Charmm36mW (with salt) which are close to those reported by Krupa *et al.*^28^ for Charmm36 and Charmm36m force fields and are considerably higher than values obtained with other Amber force fields. ^28^ One should note that the Amber99SB*-ILDN from our work is combined with the TIP4P-D water model which can lead to less compact conformations.

**Table T1:** Average values and standard deviations for the radius of gyration, end to end distance, and SASA for the three cases.

|  | Radius of gyration [nm] | EE distance [nm] | SASA [nm<sup>2</sup>] |
| --- | --- | --- | --- |
| Amber99SB*-ILDN | $1.6 \pm 0.3$ | $3.2 \pm 0.9$ | $48.9 \pm 4.7$ |
| Charmm36mW | $1.4 \pm 0.3$ | $2.7 \pm 0.9$ | $44.9 \pm 4.6$ |
| Charmm36mW with NaCl | $1.4 \pm 0.3$ | $2.6 \pm 1.0$ | $44.3 \pm 5.4$ |

These results confirm the random coil behavior of the A*β*42 monomer, dominated by constant structural fluctuations but also by the presence of extended secondary structure elements especially in the case of the Charmm36mW force field. More details regarding the recurring contacts within the monomer conformations are discussed in the context of contact maps below.

#### Contact maps and clustering

The contact maps of the three studied cases reveal intramolecular interactions that are highly related to the *β*-sheet content discussed above. Here we discuss the main features observed in the contact maps with probabilities close to or above 20%. For a better understanding of the contact patterns we also discuss at this stage the clustering of the conformations from each simulation. We show representative structures for the top five clusters with the highest populations and contact maps derived from all the conformations that belong to each cluster. The populations of the clusters are displayed in Table ST1.

In the complete contact map of Amber99SB*-ILDN from Figure S6 Left there are two main regions of contact: R5-H6 with G9-V12 and Q15-A21 with M35-G29. This first region contains contacts which have the highest probability in this simulation, i.e. H6 with G9. The second region contains a linear pattern perpendicular to the main diagonal, which is common to anti-parallel *β*-sheets and has probabilities between 20 and 30%. Given the high propensity for *β*-sheet structure in this region, 20-30% in Figure 1 Bottom, it is likely that many conformations have an anti-parallel *β*-sheet between these amino acids, i.e. Q15-A21 and M35-G29. This pattern can also be identified in the top five clusters from Figure S7 Left, where the anti-parallel *β*-sheet pattern discussed above is present in four out of five cases, the exception being cluster two. One should note that the total population of the top five clusters is 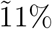, as shown in Table ST1, which means that the conformations in the full trajectory are very diverse leading to many clusters with small populations. Nevertheless, the representative structures of the top five clusters are still good examples for how the contact patterns observed in the contact maps translate into visual structures.

The simulation with the Charmm36mW force field without NaCl has in the complete contact map from Figure S6 Middle three main contact patterns with clear shapes that also correspond to anti-parallel *β*-sheets: 1) D1-G9 with F21-H14 with contact probability slightly below 20%, 2) D7-H13 with E22-G15 with probability between 20 and 30%, and H14-V24 with G38-S26 with probabilities between 30 and 60%. Part of pattern one which has the lowest probability can be identified in the contact map of cluster five from Figure S7 as a short anti-parallel *β*-sheet between D1-F4 and F19-K16 also present in the representative conformation. Pattern two is partially present in the contact maps of clusters one and two and visible in the 3D structure. The third anti-parallel *β*-sheet pattern can be clearly seen in the third and fourth clusters, and especially in the representative structure of cluster three. In addition, short patterns common to parallel *β*-sheet conformations can also be identified in the complete contact map at A2-H14 with S26-V39. These patterns are very clear in cluster two at R5-S8 with G29-I32, and cluster five at S8-H14 with I32-V39. To confirm these observations we have identified conformations from clusters two and five where these parallel *β*-sheets occur and we show them in Figure S8. Of great interest is the conformation from cluster five with the short parallel and anti-parallel *β*-sheets due to the compact or disk-like shape of the monomer. This type of structures, stabilized by parallel *β*-sheets can have very interesting characteristics as discussed below. In contrast to the Amber99SB*-ILDN simulation, for Charmm36mW the top five clusters contain 27% of the total population of conformations, almost equally distributed between the first four clusters as seen in Table ST1.

The contact map of Chamm36mW with NaCl is to some degree very similar to the Charmm36mW simulation without salt, see Figure S6. Contact patterns two and three corresponding to anti-parallel *β*-sheet structures seen in the simulation without salt are also present here, but we label them as pattern one (D7-H13 with E22-G15) and two (Y10-V24 and A42-Q25) which extends along more amino acids than the pattern from the simulation without NaCl. Pattern two is seen clearly in clusters three and four in Figure S7 Right, while pattern one is present in clusters one and five. Two contact patterns that include parallel *β*-sheets are present in this simulation at D1-Y10 with K28-G37 as pattern three and at Q15-A21 with G37-A42 as pattern four in Figure S6 and are very clearly exemplified in cluster one from Figure S7. This cluster, which contains 14% of the total population of conformations has a very special type of structure, which is discussed in detail in the following sections. This is a very significant population, given that the top five clusters amount to 28% population. Many of the features observed in the simulation with NaCl are similar with the ones observed in the simulation without salt. This is expected to some degree since both simulations use the same force field and water model. Nevertheless, the additional 150 mM NaCl did have an effect on the conformations, especially on the contacts between charged amino acids. We have calculated contact probabilities for amino acid pairs with opposite charges and we show them in Figure S9. While the mean contact probability was ~12% for both cases, there are significant differences at the level of individual contacts. The strongest contacts, for pairs K28–E22 and K28–D23, do not differ much between the two simulations. The other pairs however, with contacts below 15% probability, undergo changes when NaCl is added. Thus, interactions at D1–E22, R5–E22 and K28–E3 in the simulation without salt are reduced in amplitude, while interactions at D1–D23, K16–D42, K28–D1 and K28–A42 in the simulation with NaCl are increased in amplitude. The high contact probability for K28– E22 and K28–D23 observed here correlates well with results from previous computational studies. ^29–31^ In the case of the simulation with NaCl, the increased interactions for amino acid pairs D1–D23 and K28–A42 are most likely related to the cluster one discussed above, where these residue pairs form salt-bridges for a considerable number of conformations in the cluster.

Similar conformations to those observed here and containing anti-parallel *β*-sheets have also been observed in previous computational studies of the A*β*42 peptide. ^28,32–34^ Other studies indicate a lack of any significant extent of secondary structure in A*β*42 and relate this to either the sampling method used^20^ or the force field.^19^ We would like to highlight that the conformational diversity observed under all three simulation conditions, including the presence of conformations with a significant population of *β*-sheets, leads to a good agreement between the calculated and experimental NMR J-couplings, especially for the simulation with NaCl. These conclusions confirm that our simulations sample conformational ensembles with random coil behavior as predicted by experiments. The unique feature of molecular dynamics simulations, however, is the possibility to uncover individual conformational substates that contribute to the random coil, such as cluster one from the simulation with the Charmm36mW force field and NaCl. This unprecedented type of conformation gains its relevance from its similarity with the fibrillar state into which the peptide can convert, and is therefore discussed in detail below.

### Fibril-like A*β*42 monomer conformation

Monomeric A*β*42 conformations that bear any resemblance with peptides from fibril models have been postulated and long been sought for without much success thus far. ^26,35^ Such a conformation would have strong implications for the aggregation process, but especially for the fibril elongation by monomer addition observed in experimental studies, ^36^ because it may provide mechanistic insight into the misfolding process.

In this context, of particular interest is the cluster with the largest population from the simulation with the Charmm36mW force field and NaCl. The representative conformation is shown in Figure 2.

**Figure 2:**
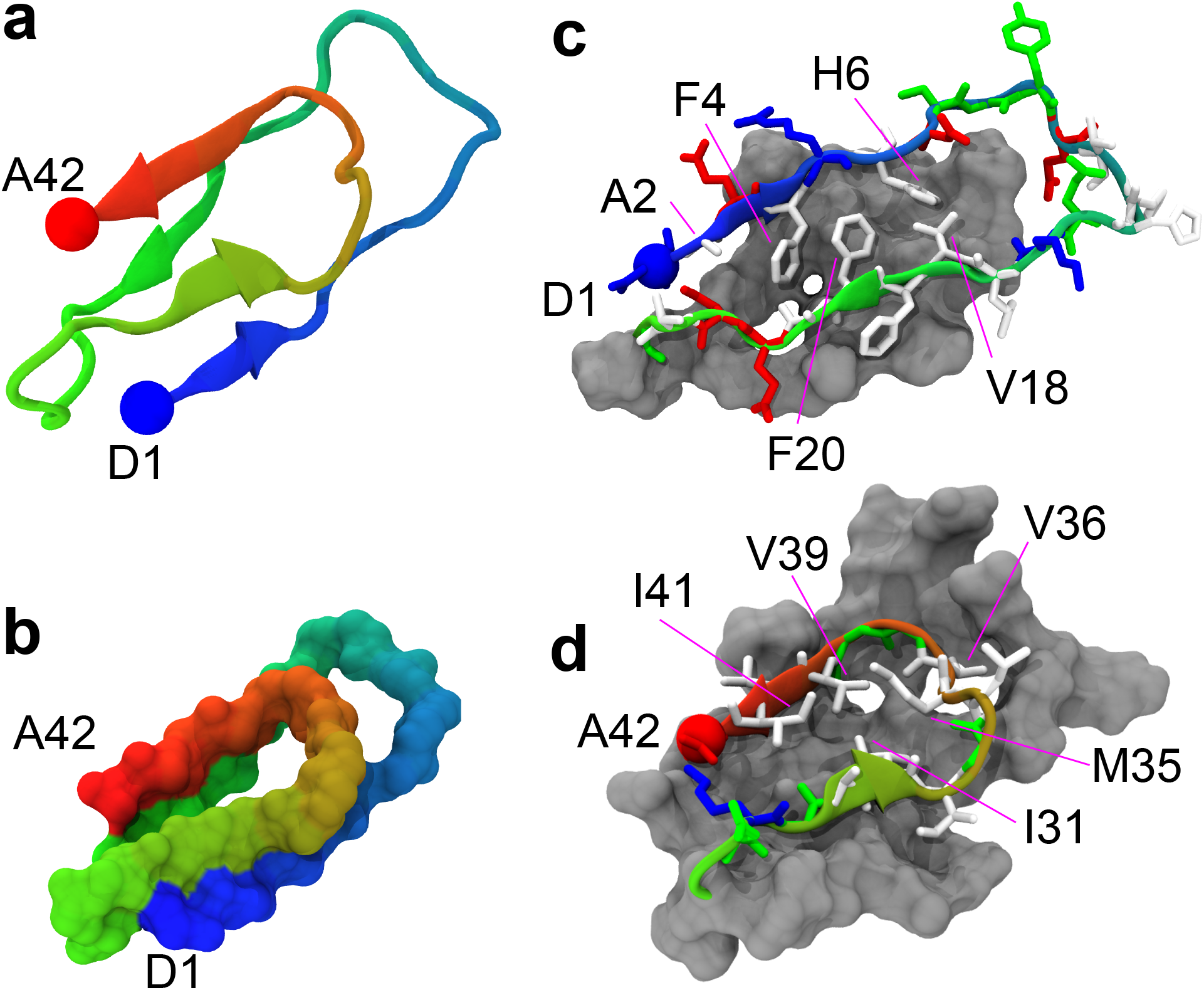
Structure of the fibril-like monomer. a) Cartoon representation highlighting the secondary structure, the N-terminus shown in blue and the C-terminus shown in red. b) van der Waals representation of the backbone which emphasizes the coiled geometry. c) N-terminus loop with all amino acids shown in licorice representation and colored by the type of amino acid (white - hydrophobic, red - negatively charged, blue - positively charged and green - polar. The C-terminus loop is shown as gray surface. d) C-terminus loop is displayed similarly to c) and the N-terminus loop is shown as gray surface.

#### Stability factors

The first defining characteristics of this structure, as seen in Figure 2a, is its compact shape and two short parallel *β*-sheets. The backbone atoms of the peptide form a spiral-like structure, Figure 2b, where the N-terminus shown in blue forms a first loop/ring until the middle part of the peptide, i.e. residue N27, comes in contact with the N-terminus. We label this region as the N-terminus loop. The rest of the peptide, K28–A42, forms a second loop which we label as the C-terminus loop. The two loops appear as two different flat surfaces stacked on top of each other. This is more evident from Figure 2c and d, where the N-terminal loop and the C-terminal loop are drawn as surfaces. Another important characteristic is the location of many hydrophobic amino acids in the interior of the two loops and shielded from the solvent to some degree. Within the N-terminal loop one can identify in Figure 2c the following hydrophobic amino acids pointing towards the interior of the loop: A2, F4, H6, V18, and F20. Within the C-terminal loop, Figure 2d, the following hydrophobic amino acids are oriented inwards: I31, M35, V36 (points toward the N-terminal loop), V39, and I41. As will be discussed below, the orientation of the amino acids within the C-terminal loop is resembling to a large degree that from the fibrillar states of the peptide. Two crucial factors that contribute to the stability of this conformation are salt bridges and parallel *β*-sheets.

The two backbone loops are both closed by salt-bridges. The N-terminal loop is stabilized by a salt-bridge between D1 and D23, Figure 3a, and the C-terminal loop is stabilized by two salt bridges formed by K28 with D23 or with A42, Figure 3b. Two of these salt bridges are possible due to the terminal capping, NH3+ for N-terminus and COO- for the C-terminus. To quantify the occurrence of the salt-bridges we have calculated distances between the two oxygen atoms of the negatively charged amino acids and the three hydrogen atoms of the positively charged ones involved in the salt bridge. The results for the three salt bridges obtained from all the conformations in cluster one of the simulation with Charmm36mW and NaCl are shown in Figures S10 (D23-D1), S11 (E22-K28), and S12 (A42-K28). In the case of salt-bridge D23-D1, most of the conformations have either the distance between oxygen O1 or oxygen O2 of residue D23 and the three hydrogens of residue D1 below 0.4 nm which is the threshold distance to qualify as a salt-bridge. This is also shown in the normalized distributions from Figure S10 Bottom where the largest peaks below 0.4 nm belong to distances between O2 and H1 or H2. The dynamics of this salt bridge is clear from the plots in S10 Middle for both a) and b), where the three hydrogens alternate as the closest atom to the oxygen O1 or O2, with oxygen O2 being preferred for shorter distances. A similar analysis is shown in Figure S11 for salt-bridge E22–K28. This salt-bridge appears in a large number of structures from cluster one, but is not as stable as D23–D1. This can be seen in Figure S11 Top and Middle for both O1 and O2. The distance distributions from Figure S11 Bottom shows a main peak at 0.31 nm followed by a smaller peak and a shoulder at values above 0.4 nm. A similar situation occurs for salt-bridge A42–K28 shown in Figure S12, where the peak corresponding to salt-bridges is the main one, but is followed by another peak at large values and has slightly lower amplitude than E22-K28.

**Figure 3:**
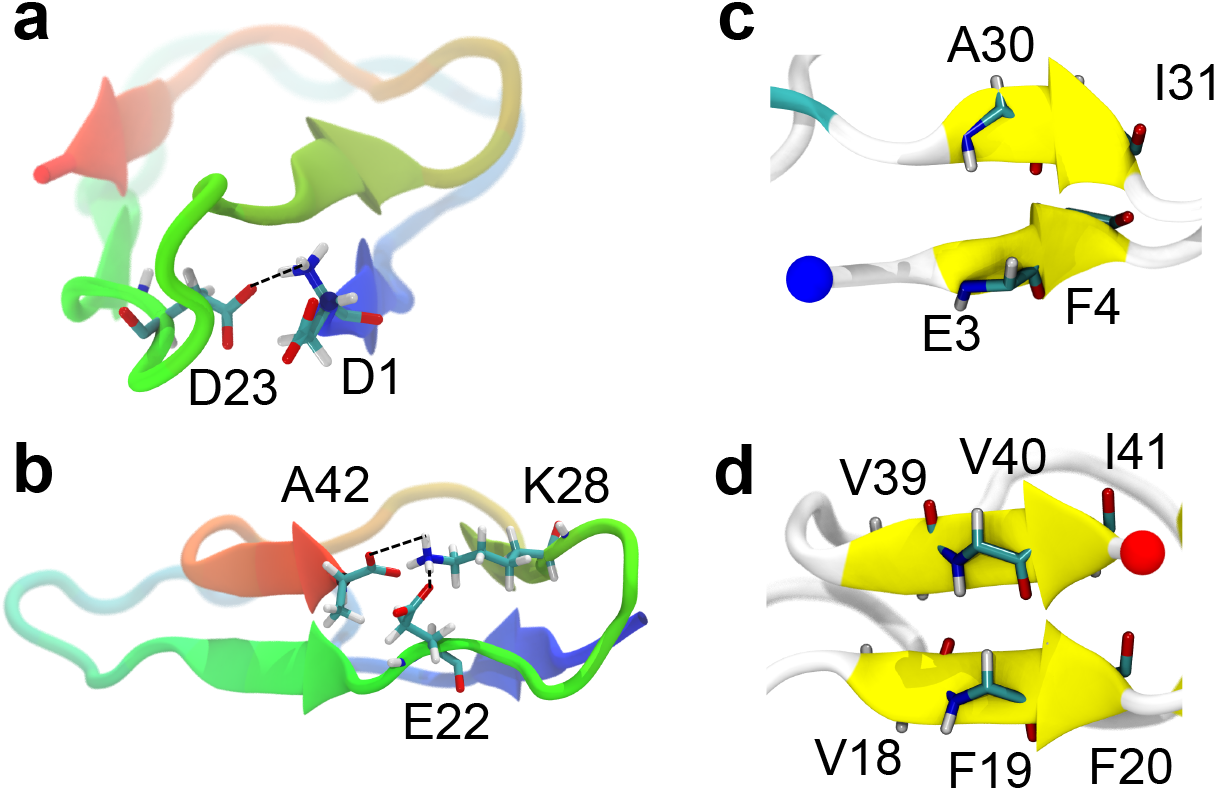
Salt-bridges and *β*-sheets of the compact monomer. a) and b) show salt-bridges D1-D23 and K28-E22/A42, respectively. c) and d) show the two parallel *β*-sheets located at E3-F4 with A30-I31 and V18-F20 with V39-I41, respectively.

Considering a salt-bridge as formed when any of the distances discussed above (between O1, O2 and H1, H2, or H3) is below 0.4 nm, we have estimated that, for this cluster, salt-bridge D23–D1 was present during 93.16% of the time the molecule spent in the fibril-like conformation, while salt-bridge E22–K28 and A42–K28 during 60.20% and 46.12% of the time, respectively. From Figure 2 it is clear that E22 and A42 are close to each other and form alternating or simultaneous salt-bridges with K28. We have identified the conformations where K28 forms salt-bridges with both E22 and A41 and calculated a propensity of 29.85%. This means that within cluster one of the simulation with NaCl, K28 was engaged in salt-bridges with either E22 or A42 in 76.47% of the conformations. Thus, both the C- and N-terminal loops are closed by salt-bridges which help to maintain the hydrophobic amino acids shielded from the solvent. Previous computational studies have identified the E22/D23-K28 salt-bridge, ^29–31^ but not the other two observed in the fibril-like structure from this study. A very recent study^37^ does mention close contacts between K28 and A42 in some of the A*β*42 monomer structures, but without the additional salt-bridges at D23–D1 and E22-K28 or the compact conformation of the N-terminus observed in our work.

In addition to the salt-bridges, two parallel *β*-sheets located at opposite positions on the spiraled conformation give further stability to this structure. The two *β*-sheets are shown in Figure 3c and d, and are formed between E3-F4 and A30-I31 in the first *β*-sheet and between V18-F20 and V39-I41 in the second *β*-sheet. They can also be identified in the contact map of cluster one from the simulation with Charmm36mW and NaCl in Figure S7 as some of the strongest contacts. One should also note other strong contacts within this cluster that do not belong to the above discussed *β*-sheets, but do contribute to the structural stability, i.e. the contacts discussed in the Contact maps and clustering section: D1–Y10 with K28–G37 and Q15–A21 with G37–A42 excluding the parallel *β*-sheets amino acids.

We have thus identified three main features of this compact structure that make it unusually stable compared to A*β*42 monomers from the other simulating conditions or from monomers reported by other studies.^19,20,28^ The salt-bridges together with the parallel *β*-sheets preserve the two big loops in contact while allowing the hydrophobic amino acids to interact with each other inside the spiraling conformation and shielded from the solvent. A recent computational study ^38^ has shown that compact conformational states, including those of monomers, are favoured during the A*β*42 assembly process and might be able to explain the oligomer size distributions observed experimentally. ^39,40^ The monomer conformation described above fits very well in this category of compact intermediates and could also contribute to the formation of compact meta-stable dimers or higher oligomers.

#### Mechanism of formation of the fibril-like structure

An important question regarding the compact, fibril-like monomer is related to the transition from an unstructured peptide into the fibril-like conformation described above. To address it we have studied the structural conversion that occurs during the H-REMD simulation. The H-REMD method generates an unbiased non-continuous trajectory, with conformations that also originate in the biased simulations. Considering all replica simulations, we identified a trajectory that contains the compact/folded structure and “jumps” between different biased simulations but is continuous in time. We have then calculated the RMSD of the backbone atoms with the meta-stable structure, i.e. center of the cluster one from the Charmm36mW simulation with NaCl, as well as the fraction of native contacts, Q, as defined by Best et al. ^41^ These quantities are shown in Figure 4 and describe a simulation interval that starts just before the onset of the folding process and ends with the occurrence of the fibril-like structure which contains all the stability elements described above. As can be seen in Figure 4 Top, the RMSD starts from a large value above 1 nm and drops gradually until it reaches a value of 0 corresponding to the structured state. Equivalently, the fraction of “native” contacts starts from a value close to 0 and increases gradually to 1, where all the contacts of the fibril-like state are formed. By analyzing these two quantities and visually inspecting this time-continuous trajectory we have identified several steps/intervals that seem crucial for the formation of the compact conformation. These steps are discussed in detail below.

**Figure 4:**
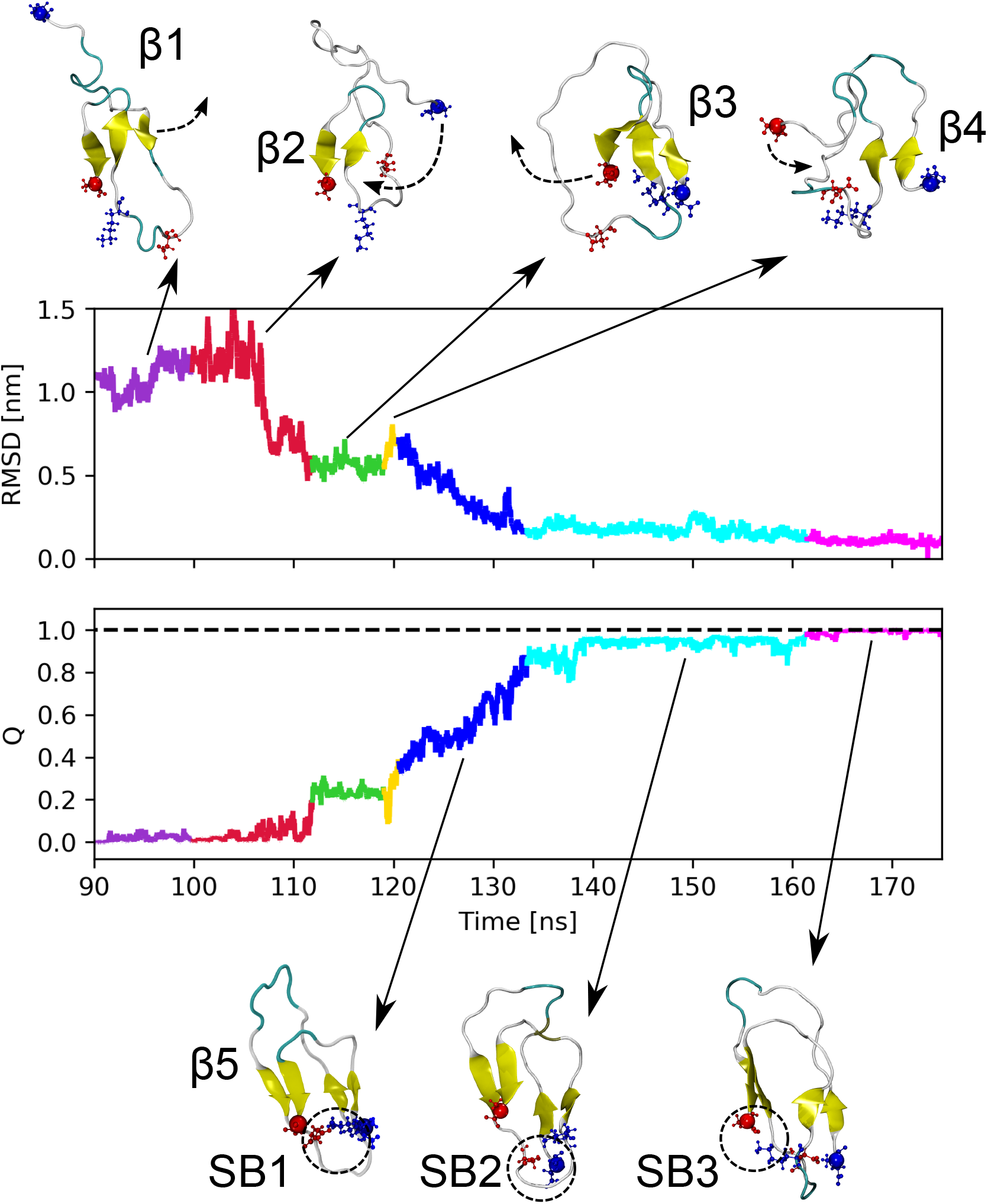
Folding of A*β*42 monomer into the compact structure. Top - RMSD of the backbone atoms between the representative monomer and each structure in the continuous time trajectory. Bottom - Fraction of native contacts Q^41^ during the continuous time trajectory. The time interval starts just before the onset of the folding and ends just after the occurrence of the representative conformation. Representative snapshots for each time interval are showed colored by secondary structure with the N and C-termini as blue and red spheres.

1) The first step shown in purple in Figure 4 corresponds to a structure which has three anti-parallel *β*-sheets (*β*1) and the N-terminus is detached from the rest of the structure. This type of structure has also been observed in cluster five. It corresponds to a large RMSD value and a low Q value. 2) The next step shown in red is the breakage of the triple *β*-sheet that results in a single anti-parallel *β*-sheet (*β*2). During this step also occurs the attachment of the N-terminus to the anti-parallel *β*-sheet. This process can be identified as a drop in the RMSD to values close to 0.6 nm and is indicated by a black dashed arrow in the second snapshot from Figure 4. 3) The resulting structure shown in the third snapshot has a triple *β*-sheet (*β*3) where the N-terminus forms a parallel sheet with the middle strand and the C-terminus forms an anti-parallel sheet with the middle strand. The rest of the peptide forms a very large loop around this *β*-sheet. This step is shown in green on the plots and corresponds to a slightly higher Q value of ~0.25. 4) The fourth step, in yellow, is the breakage of the anti-parallel *β*-sheet and the detachment of the C-terminus from the rest of the peptide. 5) From this point, the C-terminus forms gradually more and more contacts with the free loop region, outside the parallel *β*-sheet, until the second parallel *β*-sheet (*β*6) is formed between the C-terminus and the V18-F20 fragment. This process is indicated by a gradually decreasing and increasing value of the RMSD and Q, respectively, and is shown in blue in Figure 4. At this stage also forms the first salt-bridge (SB1) between E22 and K28. 6) Once the two parallel *β*-sheets are formed, the RMSD has dropped to ~0.16 nm, while the Q value has increased to ~ 0.95. This is shown in light blue in Figure 4 and during this step the second salt-bridge (SB2) between D1 and D23 forms. 7) The final step, shown in magenta, that leads to the folded conformation is the formation of the third salt-bridge (SB3) between K28 and A42 which alternates with salt-bridge K28-E22.

As described above, this is a complex structural conversion that involves large conformational changes with formation and breakage of *β*-sheets and salt-bridges. In order to obtain a more realistic estimation of the lifetime of the fibril-like conformation we have performed five classical MD simulations starting from the center of cluster one of the Charmm36mW with NaCl simulation. By considering as RMSD cutoff a value of 0.4 nm we identified an average lifetime of the conformation of 249.6 ± 41.1 ns. This is in agreement with a recent computational study which identifies metastable states of the A*β*42 monomer with sub-microsecond lifetimes.^42^ Given the fact that this type of structure appears in the unbiased simulation 14% of the time and has a considerable lifetime, it is likely that the process of structure formation described above also occurs in unbiased and continuous time simulations or in real systems, albeit at longer time scales.

#### Similarity between the fibril-like monomer and A*β*42 fibrils

One of the main features of the compact A*β*42 monomer discussed above is the structure of the middle and C-terminal region which forms a closed loop via the salt-bridge between amino acids K28 and negatively charged amino acids E22 and A42. This peptide sequence, K28-A42, has a structure very similar to peptides found in recent experimental fibril models, referred to as the S-shape fibrils. To clarify this we have aligned the structure containing the sequence K28-A42 from the compact monomer and that of fibrillar peptides from three different models as shown in Figure 5: a) a single filament fibril obtained at pH of 7.4 by Xiao *et al.*,^43^ b) a double filament fibril obtained at pH of 8 by Colvin *et al.*,^44^ and c) a double filament obtained at pH of 2 by Gremer *et al.*^45^ Note that the simulations performed in our study correspond to a pH of 7.4. The similarity between the monomer conformation obtained in our study and a peptide from each of the fibril models is quantified by the RMSD value of the backbone atoms of residues K28-A42. The smallest RMSD is obtained for the single filament fibril model of Xiao *et al.*,^43^ Figure 5a, with an RMSD of 0.15 nm for the 7th frame from the deposited PDB file with ID 2MXU. The other nine deposited frames have slightly higher RMSDs with a maximum of 0.26 nm. One important aspect of the structural comparison, visualized in Figure 5a, is the orientation of the side chains of hydrophobic amino acids from the simulated structure, which point all in the same direction as the ones from the fibrillar peptide. The same applies to K28 and A42, involved in a salt-bridge. However, in the fibril, K28 is located at a position that allows salt-bridges with A42 from the same peptide chain but also from the next peptide chain within the fibril. In the case of the double filament structure published by Colvin et al., ^44^ the smallest RMSD has a value of 0.19 nm for frame 9 from the structure with PDB ID 5KK3, and a maximum value of 0.21 nm. Figure 5b shows the case with the smallest RMSD where all hydrophobic amino acids point in the same direction, except for M35. We also compared the folded monomer structure with a peptide from the fibril obtained at pH of 2 by Gremer et al. ^45^ In this case, Figure 5c, the RMSD with the single deposited PDB structure with ID 5OQV has a value of 0.17 nm. Despite the very small backbone RMSD, only L34 and M35 from the monomer point in the same direction as the residues in the fibrillar peptide, while I31, V39 and I41 point toward the exterior of the C-terminal loop in the fibril peptide and towards the interior of the loop in the simulated peptide. These mismatches are not surprising, given the effect of the low pH on the protonation state of charged amino acids and eventually on the overall structure of the fibril.

**Figure 5:**
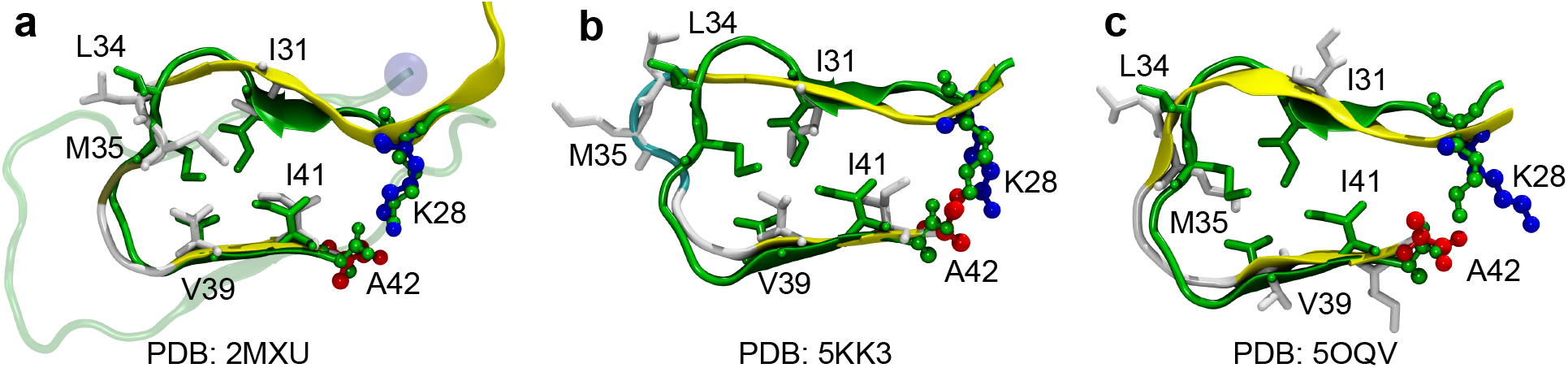
Comparison of the compact monomer with S-shape fibril models. All peptides are shown in cartoon representation and selected amino acids as licorice or balls and sticks. The fibril-like monomer state is colored in green while the fibrillar peptide is colored based on its secondary structure, its hydrophobic amino acids in white, and the charged residues K28 in blue and A42 in red. Three fibril models were considered with PDB IDs: a) 2MXU, ^43^ b) 5KK3,^44^ and c) 5OQV.^45^

As shown above, the C-terminal loop of the folded conformation is in very good agreement with all three fibril models. This means that the observed A*β*42 monomer is able to sample, besides a multitude of disordered states, partly structured conformations that mimic the A*β*42 fibril core without interfering with the overall random coil behavior of the conformational ensemble. A recent computational study using a coarse grained protein model^37^ reports a slightly similar C-terminal loop region with a close contact between K28 and A42, but with a disordered N-terminus. This study ^37^ provides evidence that although not all, some of the characteristics of the compact monomer from our work are also present under other simulation conditions. The high similarity between the C-terminal loop of the compact monomer from our work and fibrillar peptides is thus quite robust and highly relevant due to its stability provided by the N-terminus loop conformation. In fact, without the interactions with the N-terminal loop, the fibril-like features of the C-terminal loop would probably have a much lower stability and therefore shorter lifetime.

#### The fibril-like monomer optimally fits only at one of the two fibril ends

The folded monomer conformation observed in our simulations and discussed above has many characteristics similar to the fibril core. In addition, the N-terminal loop is covering completely one side of the C-terminal loop, see Figure 2. We have investigated how such a conformation could fit at the end of a fibril assuming an in-register interaction between the C-terminal loop of the folded monomer and the C-terminal region of a peptide at the fibril end. To identify which end of the fibril is optimal for this type of docking, we have aligned the folded monomer with a peptide from the fibril end using only the C-terminal loop (amino acids K28-A42) backbone atoms, and making sure that the N-terminal loop is located towards the solvent and not towards the fibril. It is important to note that we assume here that the conformation of the fibril end is identical to the overall fibrillar conformation. The structural fibril models used here correspond to the average arrangements of monomers inside the fibrils, whereas it can be expected that the fibril ends undergo some degree of structural rearrangement, given the smaller number of interaction partners compared to a monomer deeper inside the fibril. The result of these docking experiments can be seen in Figure 6 from two different angles. From Figure 6a it is clear that the folded monomer has a near-optimal alignment with the even tip of the fibril which has the C-terminus less exposed to the solvent.

**Figure 6:**
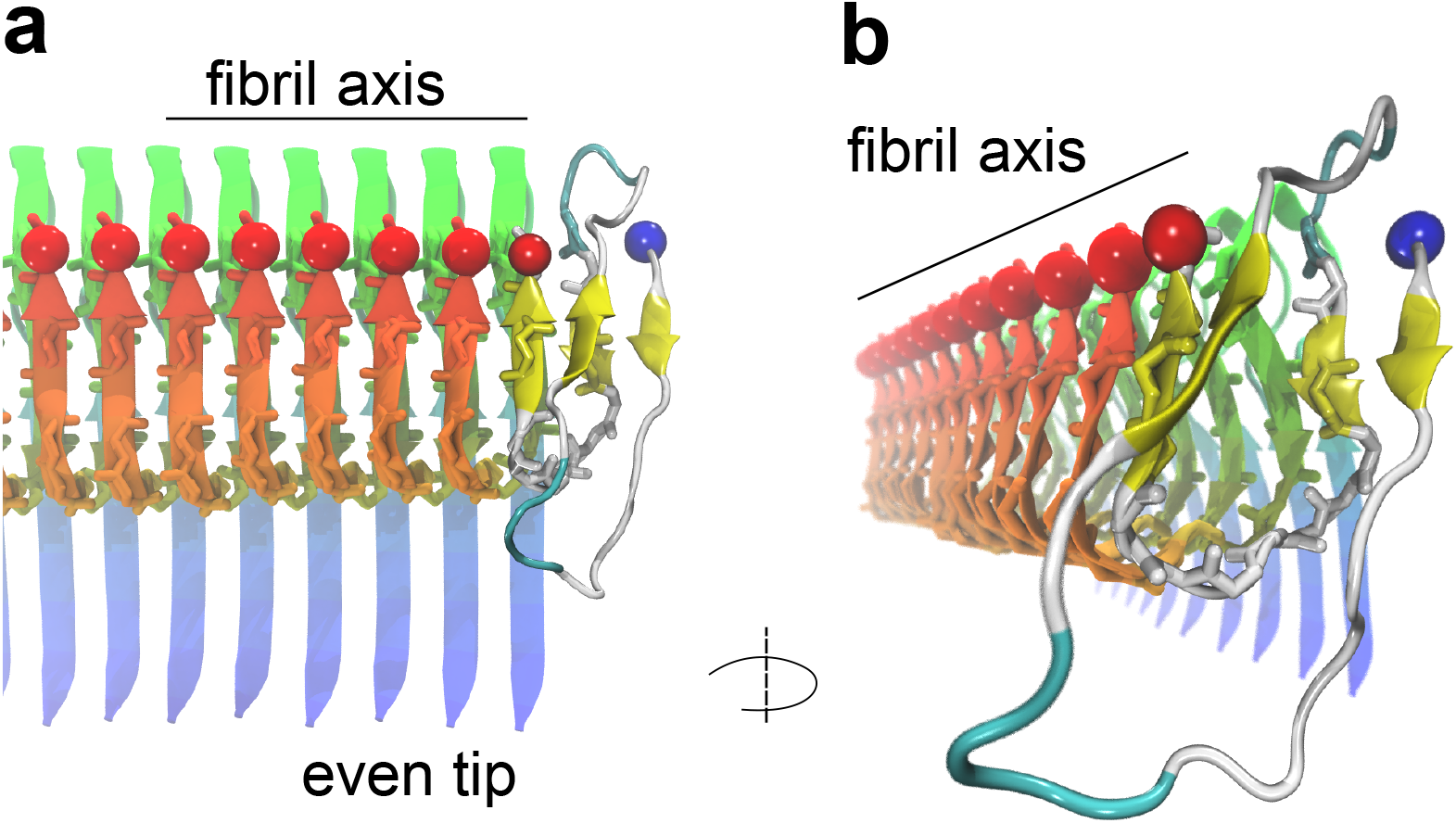
Structural fit of the compact fibril-like monomer at the fibril end. a) The fibril-like monomer observed in our work is shown at the even fibril end (the C-terminus less exposed to the solvent than the middle and N-terminal regions). All peptides are shown in cartoon representations with the simulated monomer colored by the secondary structure elements and the fibrillar peptides with color gradient from blue at N-terminus to red at C-terminus. A licorice representation of the backbone is also included to emphasize the hydrogen bonds between the peptides. The N- and C-termini are shown as blue and red spheres, respectively. b) The same structure from a) is slightly rotated for a better view along the fibril axis.

These results are interesting in the context of the discussion of two proposed mechanisms for fibril elongation: “fast deposition” or “direct docking” and “dock-lock”. During the “fast-deposition”, a so-called activated monomer, i.e. a monomer with appropriate conformation, is efficiently deposited onto a fibril end and thus becomes incorporated into the fibril.^46,47^ The compact monomer from our study would most likely fulfill the structural requirements for an activated conformation involved in the fast deposition process. However, the direct docking mechanism has been shown to contribute less to the fibril elongation process compared to the two phase “dock-lock” mechanism.^48^ In “dock-lock”, during the “dock” phase a monomer attaches somewhat non-specifically and therefore rather weakly to the fibril end as well as possibly also the fibril surface in a fully reversible manner, ^36^ while during the “lock” phase, which takes place on a much slower time scale than the “dock” phase,^49^ the monomer undergoes a structural rearrangement and samples the different *β*-sheet registers until it either dissociates off again or it achieves the most stable, correctly incorporate state, which often for practical purposes appears to be irreversible. This mechanism of fibril elongation has been proposed from experiments^50^ and was also observed in several computational studies, ^47,48^ with a clear prevalence over the the fast deposition mechanism.

Estimates of the residency time from experiments, ^36^ i.e. the time that a monomer spends on average in proximity of the fibril end, indicate that this time is several orders of magnitude larger than the lifetime of the fibril-like state estimated from our simulations. This would indicate that the monomer undergoes dozens to hundreds of transitions from fully random coil to the compact, fibril-like state during the time scale of its average contact with the fibril end. The lifetime of the fibril like state observed in our simulations of the monomer could of course also be significantly larger in close proximity to the fibril end. Regardless of whether or not the fibril-like state actually is on the pathway to an incorporated monomer, the identification of such a class of fibril-like structures allows us to rationalise certain features of the aggregation behaviour of the A*β* peptide. It has been consistently found that the A*β*42 peptide has the smallest known free energy barrier for fibril elongation of all amyloid polypeptides. ^36,51,52^ This could be explained by the innate tendency of the A*β* sequence to adopt conformations similar to those inside the fibril; given that even the isolated monomer has already a significant tendency to adopt fibril-like structures, the presence of a templating fibril end can render the conversion into the fibril-like conformation even easier, ultimately resulting in a free energy barrier of only a few k_*B*_T. The importance of the fibril-like state that the monomer adopts in isolation for the elongation reaction depends on whether the templating fibril end should be regarded as a strong or weak perturbation of the monomolecular free energy landscape explored in the present work. Some experimental evidence for an important role of the fibril end comes from the observation that the two ends of a fibril often grow with different rates,^53^ in agreement with the different efficiency in docking to the two ends, as demonstrated in this work. However, answering this question will ultimately require more extensive simulations of monomer-fibril interactions. Our contribution to this ongoing discussion is one of the most detailed characterisations of the monomolecular free energy landscape of the A*β* monomer, which has allowed us to identify the fibril-like state and which can be used as a benchmark with which to compare the monomer in contact with a fibril end.

## Conclusions

Using extensive enhanced molecular dynamics simulations, we have shown that the A*β*42 monomer adopts a random coil structure when employing different force fields and simulation conditions. We further discovered a transient monomer structure that accounts for 14% population of the total observed conformations, when using the Charmm36mW force field with 150 mM NaCl. This structure, stabilized by short parallel *β*-sheets and three salt-bridges, has many characteristics similar to fibrillar forms of the A*β*42 peptide. Specifically, the structure of the C-terminal region from K28 to A42 is very similar to the structure of the same region from the so-called S-shape fibrils, including the orientation of side chains of hydrophobic amino acids. To the best of our knowledge this is the first A*β*42 monomeric structural ensemble to feature a state with such a high degree of similarity with the fibril core. Despite its stability, manifest through sub-microsecond lifetimes, this type of monomer does not alter the overall random coil behavior of the structural ensemble. This suggests that the A*β*42 peptide in monomeric form already has an affinity to sample structural features specific to fibrils and could explain why A*β*42 has the lowest free energy barrier for activating the fibril elongation among amyloidogenic proteins. ^51^ Further investigation is needed to identify how exactly is this property of the compact A*β*42 monomer involved in the elongation process. In our study, the compact monomer state has the C-terminal loop, residues K28–A42, in a conformation similar to that of fibril core, which might be an indication that this region is indeed the first to form when a peptide locks onto an existing fibril end. It will be important to investigate in future studies whether this conformation also appears in the proximity of another monomer or the fibril end and how exactly is it involved in molecular mechanisms of fibril elongation such as the direct docking or the “dock-lock” mechanism.

## Methods

### MD simulations details

We have studied the conformational flexibility of the A*β*42 monomer in water by performing H-REMD^54^ simulations with several force fields and water models with a cumulated simulation time of 157.4 *μ*s. The first force field - water model combination (labeled as Amber99SB*-ILDN) includes the Amber99SB*-ILDN force field^21,22^ and the TIP4P-D water model.^23^ This water model was specifically developed to accurately describe intrinsically disorder proteins (IDPs) in molecular dynamics (MD) simulations by using and increased protein-water interaction. The second combination (labeled as Charmm36m) is the Charmm36m force field ^24^ and the Charmm TIP3P water model designed for folded proteins and IDPs. In addition, we also considered the Charmm36m force field with a modified Charmm TIP3P water model that leads to an increased protein - water interaction, more similar to the TIP4P-D model, and labeled as Charmm36mW. ^24^ In Charmm36mW, the Lennard Jones well depth parameter *ϵ* of the hydrogen atoms have been modified from −0.046 kcal/mol to −0.10 kcal/mol as suggested by Huang et al. ^24^ Simulations with Charmm36mW were considered with and without 150 mM NaCl concentration in order to assess the effect of salt ions on the protein conformation.

All simulations were performed with the Gromacs 2016.04 parallel software package. ^55^ Short range electrostatics and van der Waals interactions were cut at 0.1 nm, while long range electrostatic interactions were treated with the Particle Mesh Ewald method. The temperature was kept at 300 K via velocity rescaling with a stochastic term algorithm ^56^ and a time constant for coupling of 0.1 ps. The pressure coupling was controlled with the Parrinello-Rahman barostat^57,58^ with a time constant of 1 ps. The hydrogen atoms were treated as virtual interaction sites, allowing an integration time step of 4 fs while maintaining energy conservation. ^59^

The solution structure of A*β*42 protein monomer with PDB ID 1IYT was used as starting conformation. ^7^ The conformation was placed in a dodecahedral box with 1.6 nm between the protein and the box, solvated with water molecules and 3 Na ions added for charge neutrality, except for the second simulation with the Charmm36mW force field where 43 Na and 40 Cl ions were added to also account for 150 mM salt concentration. The final systems had the following number of atoms: 1) Amber99SB*-ILDN - 62,150 atoms; Charmm36m - 47,214 atoms; Charmm36mW - 47,214; Charmm36mW with NaCl - 47,054. This simulation box was large enough to allow free translation and rotation of the A*β*42 monomer without interacting with its periodic images that would otherwise result in simulation artifacts. After energy minimization, position restrained equilibration (with the Berendsen barostat) and a short free equilibration (with the Parrinello-Rahman barostat), Hamiltonian replica exchange ^54^ simulations were was performed for each system. Each H-REMD simulation consisted of 10 (Amber99SB*-ILDN) or 12 (Charmm36m and Charmm36mW) simulations run in parallel, each simulation having a different interaction Hamiltonian where non-bonded interactions and dihedral angles are scaled with a factor *λ*. The biasing coefficients lambda can be expressed as an inverse temperature (1/temperature) correspond to temperatures between 300 and 500 K and assigned to the replicas according to a geometric distribution. The average replica exchange probability for the four systems had values between 16% and 24%.

In total we performed four H-REMD simulations, one for each system, of different length/replica: Amber99SB*-ILDN – 2 *μ*s; Charmm36m – 2 *μ*s; Charmm36mW – 3 *μ*s; Charmm36mW with NaCl – 3.4 *μ*s. All simulations were performed on the supercomputer JURECA^60^ at the Jülich Supercomputing Centre (JSC).

### Analysis

Since all simulations started from the A*β*42 structure with the PDB code ID 1IYT,^7^ the initial states have a large number of amino acids in helical conformation, see Figure S1. Except for the simulation with Charmm36m, in all other cases the highly helical conformation drops to small values below 10 residues, either within the first 100 ns for Amber99SB*-ILDN or within 1,500 ns for the simulations with the Charmm36mW force field, as can be seen in Figure S1. The drop in helical conformation is accompanied by an increase in the *β*-sheet content, especially for Charmm36mW. This is not the case for Charmm36m, where the helical residues fluctuates throughout the 2,000 ns close to a value of 30, similar to the initial state, despite a slight increase in *β*-sheet content. This indicates that the simulation takes much longer to equilibrate than for the other force fields. Given the random coil character of the A*β*42 monomer observed in experiments,^9^ we have decided to analyze only the equilibrated part of each simulation, where fluctuations in the secondary structure are similar to that of random coil, i.e. without many amino acids in helical conformation. Thus, the trajectory intervals used for the analysis consist of the last 1,000 ns from the simulations with Amber99SB*-ILDN, and Charmm36mW with and without NaCl, and are highlighted in Figure S1 with two red vertical bars.

^3^*J_HNHα_* **NMR scalar couplings** were calculated with the Karplus equation for each amino acid:

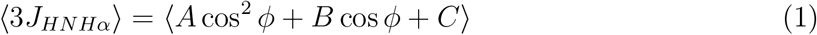

with coefficients *A* = 7.97 Hz, *B* = - 1.26 Hz and *C* = 0.63 Hz.^61^ The comparison with experimental data was done using the reduced *χ*^2^:

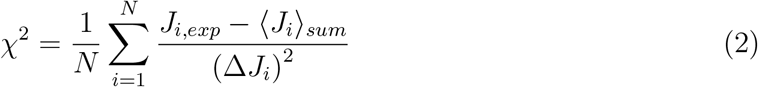

where (Δ*J_i_*)^2^ = (Δ_block_)^2^ + (Δ_Karplus_)^2^, Δ_block_ being the simulation error calculated with block averaging and Δ_Karplus_ = 0.42 Hz, the experimental error.

### Secondary structure, contact maps and clustering

Secondary structure propensities per residues were calculated with the program DSSP^62^ and the errors with block averaging. For snapshots of the protein structure we used the program Visual Molecular Dynamics (VMD)^63^ where the secondary structure is calculated with STRIDE. ^64^ Contact maps were calculated using the Contact Map Explorer module implemented in Python and considering a contact between two amino acids when any two atoms from the two residues were found at a distance below a cutoff of 0.5 nm. Clustering of structures was performed with the Daura algorithm^65^ implemented in Gromacs using the backbone atoms and an RMSD cutoff of 0.4 nm.

### Radius of gyration, hydrodynamic radius, end to end distance and SASA

The radius of gyration was calculated using the function compute-rg from the MDTraj library ^66^ implemented in Python. From the radius of gyration we calculated the hydrodynamic radius of each conformation using an empirically parametrized equation specifically derived for intrinsically disordered proteins.^27^ The end to end distance was calculated between the *C_α_* atoms of the first and last amino acids using the function compute-distances from MDTraj. To calculate the SASA we used the shrake-rupley algorithm ^67^ implemented in MDTraj.

## Acknowledgement

The authors gratefully acknowledge the computing time granted by the John von Neumann Institute for Computing (NIC) and provided on the supercomputer JURECA at Jülich Supercomputing Centre (JSC). We also acknowledge DFG grant BA 5956/2-1 for funding the current research. AKB thanks the Novo Nordisk Foundation for support through a Novo Nordisk Foundation Professorship (NNFSA170028392).

## Supporting Information Available

Additional figures and tables are included in Supporting Information.

## Notes

### Competing Interest Statement

The authors have declared no competing interest.

## References

(1) Scheuner, D. et al. Nature Med. 1996, 2, 864–870.

(2) Suzuki, N.; Cheung, T. T.; Cai, X. D.; Odaka, A.; Otvos, L.; Eckman, C.; Golde, T. E.; Younkin, S. G. Science 1994, 264, 1336–1340.

(3) Knowles, T. P. J.; Vendruscolo, M.; Dobson, C. M. Nat. Rev. Mol. Cell. Biol. 2014, 15, 384–396.

(4) Selkoe, D. J.; Selkoe, D. J. Annu. Rev. Neurosci. 1994, 17, 489–517.

(5) Esler, W. P.; Wolfe, M. S. Science 2001, 293, 1449–1454.

(6) Lichtenthaler, S. F.; Wang, R.; Grimm, H.; Uljon, S. N.; Masters, C. L.; Beyreuther, K. Proc. Natl. Acad. Sci. U.S.A. 1999, 96, 3053–3058.

(7) Crescenzi, O.; Tomaselli, S.; Guerrini, R.; Salvadori, S.; D’Ursi, A. M.; Temussi, P. A.; Picone, D. Eur. J. Biochem. 2002, 269, 5642–5648.

(8) Tomaselli, S.; Esposito, V.; Vangone, P.; van Nuland, N. A. J.; Bonvin, A. M. J. J.; Guerrini, R.; Tancredi, T.; Temussi, P. A.; Picone, D. ChemBioChem 2006, 7, 257–267.

(9) Roche, J.; Shen, Y.; Lee, J. H.; Ying, J.; Bax, A. Biochemistry 2016, 55, 762.

(10) Lim, K. H.; Collver, H. H.; Le, Y. T. H.; Nagchowdhuri, P.; Kenney, J. M.; Lim, K. H.; Collver, H. H.; Le, Y. T. H.; Nagchowdhuri, P.; Kenney, J. M. Biochem. Biophys. Res. Commun. 2007, 353, 443–449.

(11) Ono, K.; Condron, M. M.; Teplow, D. B. Proc. Natl. Acad. Sci. 2009, 106, 14745–14750.

(12) Granata, D.; Baftizadeh, F.; Habchi, J.; Galvagnion, C.; De Simone, A.; Camilloni, C.; Laio, A.; Vendruscolo, M. Sci. Rep. 2015, 5, 1–15.

(13) Nag, S.; Sarkar, B.; Bandyopadhyay, A.; Sahoo, B.; Sreenivasan, V. K. A.; Kombrabail, M.; Muralidharan, C.; Maiti, S. J. Biol. Chem. 2011, 286, 13827–13833.

(14) Mittag, J. J.; Milani, S.; Walsh, D. M.; Rädler, J. O.; McManus, J. J. Biochem. Biophys. Res. Commun. 2014, 448, 195–199.

(15) Schneider, M.; Walta, S.; Cadek, C.; Richtering, W.; Willbold, D. Sci. Rep. 2017, 7, 1–8.

(16) Wennmalm, S.; Chmyrov, V.; Widengren, J.; Tjernberg, L. Anal. Chem. 2015, 87, 11700–11705.

(17) Novo, M.; Freire, S.; Al-Soufi, W. Sci. Rep. 2018, 8, 1–8.

(18) Nasica-Labouze, J. et al. Chem. Rev. 2015, 115, 3518–3563.

(19) Meng, F.; Bellaiche, M. M. J.; Kim, J.-Y.; Zerze, G. H.; Best, R. B.; Chung, H. S. Biophys. J. 2018, 114, 870–884.

(20) Lincoff, J.; Sasmal, S.; Head-Gordon, T. J. Chem. Phys. 2019, 150, 104108.

(21) Lindorff-Larsen, K.; Piana, S.; Palmo, K.; Maragakis, P.; Klepeis, J. L.; Dror, R. O.; Shaw, D. E. Proteins 2010, 78, 1950–1958.

(22) Best, R. B.; Hummer, G. J. Phys. Chem. B 2009, 113, 9004–9015.

(23) Piana, S.; Donchev, A. G.; Robustelli, P.; Shaw, D. E. J. Phys. Chem. B 2015, 119, 5113–5123.

(24) Huang, J.; Rauscher, S.; Nawrocki, G.; Ran, T.; Feig, M.; de Groot, B. L.; Grubmller, H.; MacKerell Jr, A. D. Nat. Methods 2017, 14, 71–73.

(25) Yan, Y.; McCallum, S. A.; Wang, C. J. Am. Chem. Soc. 2008, 130, 5394–5395.

(26) Rosenman, D. J.; Connors, C. R.; Chen, W.; Wang, C.; García, A. E. J. Mol. Biol. 2013, 425, 3338–3359.

(27) Nygaard, M.; Kragelund, B. B.; Papaleo, E.; Lindorff-Larsen, K.; Nygaard, M.; Kragelund, B. B.; Papaleo, E.; Lindorff-Larsen, K. Biophys. J. 2017, 113, 550–557.

(28) Krupa, P.; Quoc Huy, P. D.; Li, M. S. J. Chem. Phys. 2019, 151, 055101.

(29) Tarus, B.; Straub, J. E.; Thirumalai, D. J. Am. Chem. Soc. 2006, 128, 16159–16168.

(30) Côté, S.; Derreumaux, P.; Mousseau, N. J. Chem. Theory Comput. 2011, 7, 2584–2592.

(31) Barz, B.; Urbanc, B. PLoS One 2012, 7, e34345.

(32) Ball, K. A.; Phillips, A.; Wemmer, D.; Head-Gordon, T. Biophys. J. 2013, 104, 2714–2724.

(33) Liao, Q.; Owen, M. C.; Olubiyi, O. O.; Barz, B.; Strodel, B. Isr. J. Chem. 2017, 57, 771–784.

(34) Carballo-Pacheco, M.; Strodel, B. Protein Sci. 2017, 26, 174–185.

(35) Frigori, R. B.; Barroso da Silva, F. L.; Carvalho, P. P. D.; Alves, N. A. J. Phys. Chem. B 2020, 124, 2798–2805.

(36) Buell, A. K. Biochem. J. 2019, 476, 2677–2703.

(37) Chakraborty, D.; Straub, J. E.; Thirumalai, D. bioRxiv 2020, DOI:2020.02.09.940676 preprint.

(38) Barz, B.; Liao, Q.; Strodel, B. J. Am. Chem. Soc. 2018, 140, 319–327.

(39) Bitan, G.; Kirkitadze, M. D.; Lomakin, A.; Vollers, S. S.; Benedek, G. B.; Teplow, D. B. Proc. Natl. Acad. Sci. 2003, 100, 330–335.

(40) Bernstein, S. L.; Dupuis, N. F.; Lazo, N. D.; Wyttenbach, T.; Condron, M. M.; Bitan, G.; Teplow, D. B.; Shea, J.-E.; Ruotolo, B. T.; Robinson, C. V.; Bowers, M. T. Nat. Chem. 2009, 1, 326–331.

(41) Best, R. B.; Hummer, G.; Eaton, W. A. Proc. Natl. Acad. Sci. U.S.A. 2013, 110, 17874–17879.

(42) Löhr, T.; Kohlhoff, K.; Heller, G. T.; Camilloni, C.; Vendruscolo, M. bioRxiv 2020, DOI:2020.05.07.082818 preprint.

(43) Xiao, Y.; Ma, B.; McElheny, D.; Parthasarathy, S.; Long, F.; Hoshi, M.; Nussinov, R.; Ishii, Y. Nat. Struct. Mol. Biol. 2015, 22, 499–505.

(44) Colvin, M. T.; Silvers, R.; Ni, Q. Z.; Can, T. V.; Sergeyev, I.; Rosay, M.; Donovan, K. J.; Michael, B.; Wall, J.; Linse, S.; Griffin, R. G. J. Am. Chem. Soc. 2016, 138, 9663–9674.

(45) Gremer, L.; Schölzel, D.; Schenk, C.; Reinartz, E.; Labahn, J.; Ravelli, R. B. G.; Tusche, M.; Lopez-Iglesias, C.; Hoyer, W.; Heise, H.; Willbold, D.; Schröder, G. F. Science 2017, 358, 116–119.

(46) Massi, F.; Straub, J. E. Proteins 2001, 42, 217–229.

(47) Straub, J. E.; Thirumalai, D. Annu. Rev. Phys. Chem. 2011, 62, 437–463.

(48) Cao, Y.; Tang, X.; Yuan, M.; Han, W. In Computational Approaches for Understanding Dynamical Systems: Protein Folding and Assembly; Strodel, B., Barz, B., Eds.; Progress in Molecular Biology and Translational Science; Academic Press, 2020; Vol. 170; pp 461–504.

(49) Buell, A. K.; Blundell, J. R.; Dobson, C. M.; Welland, M. E.; Terentjev, E. M.; Knowles, T. P. J.; Buell, A. K.; Blundell, J. R.; Dobson, C. M.; Welland, M. E.; Terentjev, E. M.; Knowles, T. P. J. Phys. Rev. Lett. 2010, 104, 228101.

(50) Esler, W. P.; Stimson, E. R.; Jennings, J. M.; Vinters, H. V.; Ghilardi, J. R.; Lee, J. P.; Mantyh, P. W.; Maggio, J. E. Biochemistry 2000, 39, 6288–6295.

(51) Buell, A. K.; Dhulesia, A.; White, D. A.; Knowles, T. P. J.; Dobson, C. M.; Welland, M. E. Angew. Chem. Int. Ed. 2012, 51, 5247–5251.

(52) Cohen, S. I. A. et al. Nat. Chem. 2018, 10, 523–531.

(53) Young, L. J.; Schierle, G. S. K.; Kaminski, C. F. Phys. Chem. Chem. Phys. 2017, 19, 27987–27996.

(54) Bussi, G. Mol. Phys. 2014, 112, 379–384.

(55) Hess, B.; Kutzner, C.; van der Spoel, D.; Lindahl, E. J. Chem. Theory. Comput. 2008, 4, 435–447.

(56) Bussi, G.; Donadio, D.; Parrinello, M. J. Chem. Phys. 2007, 126, 014101.

(57) Parrinello, M.; Rahman, A. J. Appl. Phys. 1981, 52, 7182–7190.

(58) Nosé, S.; Klein, M. L. Mol. Phys. 1983, 50, 1055–1076.

(59) Feenstra, K. A.; Hess, B.; Berendsen, H. J. C. J. Comput. Chem. 1999, 20, 786–798.

(60) Jülich Supercomputing Centre, Journal of large-scale research facilities 2018, 4.

(61) Vögeli, B.; Ying, J.; Grishaev, A.; Bax, A. J. Am. Chem. Soc. 2007, 129, 9377–9385.

(62) Kabsch, W.; Sander, C. Biopolymers 1983, 22, 2577–2637.

(63) Humphrey, W.; Dalke, A.; Schulten, K. Journal of Molecular Graphics 1996, 14, 33–38.

(64) Frishman, D.; Argos, P. Proteins: structure, function and genetics 1995, 23, 566–579.

(65) Daura, X.; Gademann, K.; Jaun, B.; Seebach, D.; van Gunsteren, W. F.; Mark, A. E. Angew. Chem. Int. Ed. 1999, 38, 236–240.

(66) McGibbon, R. T.; Beauchamp, K. A.; Harrigan, M. P.; Klein, C.; Swails, J. M.; Hernández, C. X.; Schwantes, C. R.; Wang, L.-P.; Lane, T. J.; Pande, V. S. Biophysical Journal 2015, 109, 1528–1532.

(67) Shrake, A.; Rupley, J. A. J. Mol. Biol. 1973, 79, 351–371.

