## Supporting information for "Monomeric amyloid *β*-peptide (1-42) significantly populates compact fibril-like conformations"

### Supporting information: Monomeric amyloid $\beta$-peptide (1-42) significantly populates compact fibril-like conformations

Bogdan Barz,[*,†,‡] Alexander K. Buell,[†,¶] and Soumav Nath[†,‡]

†*Institut für Physikalische Biologie, Heinrich-Heine-Universität Düsseldorf, Düsseldorf, Germany*

‡*Institute of Biological Information Processing - Structural Biochemistry (IBI-7), Research Centre Jülich, Jülich, Germany*

¶*Department of Biotechnology and Biomedicine, Technical University of Denmark, Lyngby, Denmark*

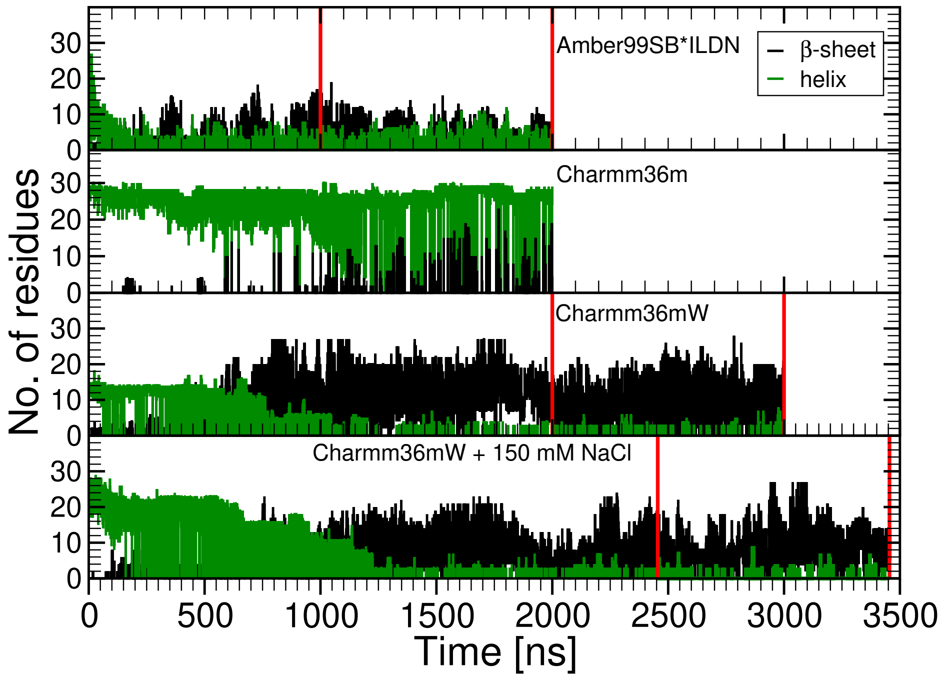


Figure S1: Evolution of the number of residues adopting sheet or helix conformations during the H-REMD simulations. The four panels display the data for the four conditions: Amber99SB*-ILDN with TIP4P water, Charmm36m with TIP3P water, Charmm36mW with TIP3P water, and Charmm36mW with TIP3P water and 150 mM NaCl.

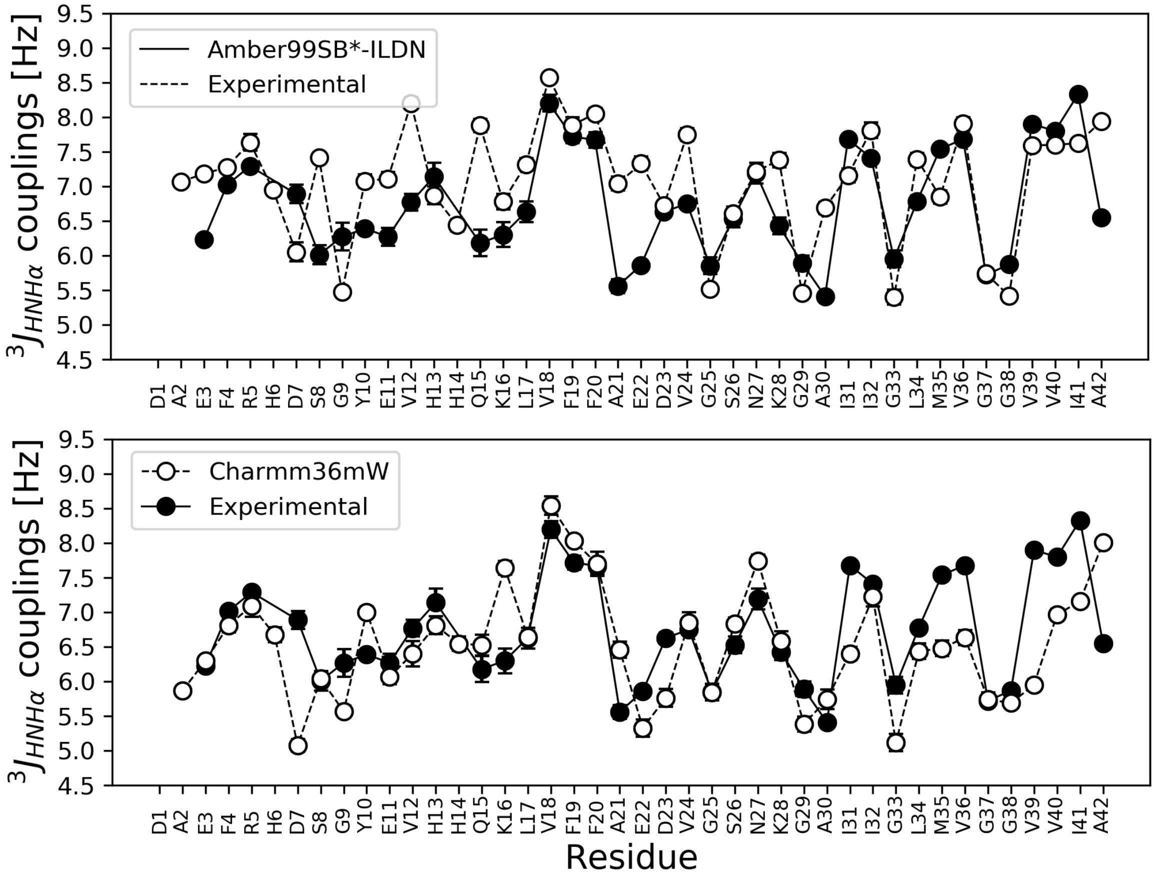


Figure S2: 3J$_{HNH\alpha}$ NMR scalar couplings per residue for the Amber99SB*-ILDN (Top) and Charmm36mW force field without NaCl (Bottom). In black are shown experimental values and in white those obtained from the H-REMD simulations.

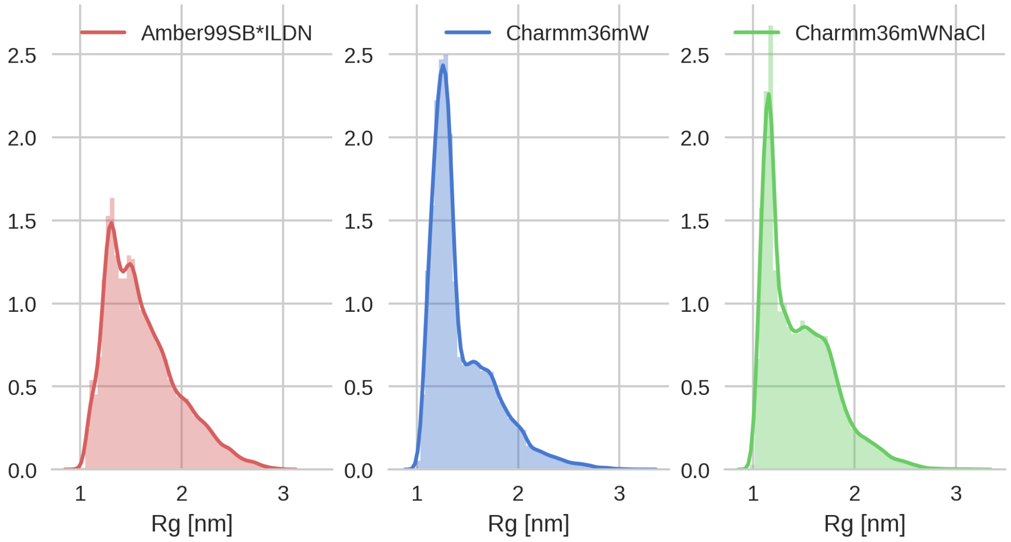


Figure S3: Radius of gyration normalised distributions for the three cases: (A) Amber99SB*-ILDN, (B) Charmm36mW, and (C) Charmm36mW with 150 mM NaCl.

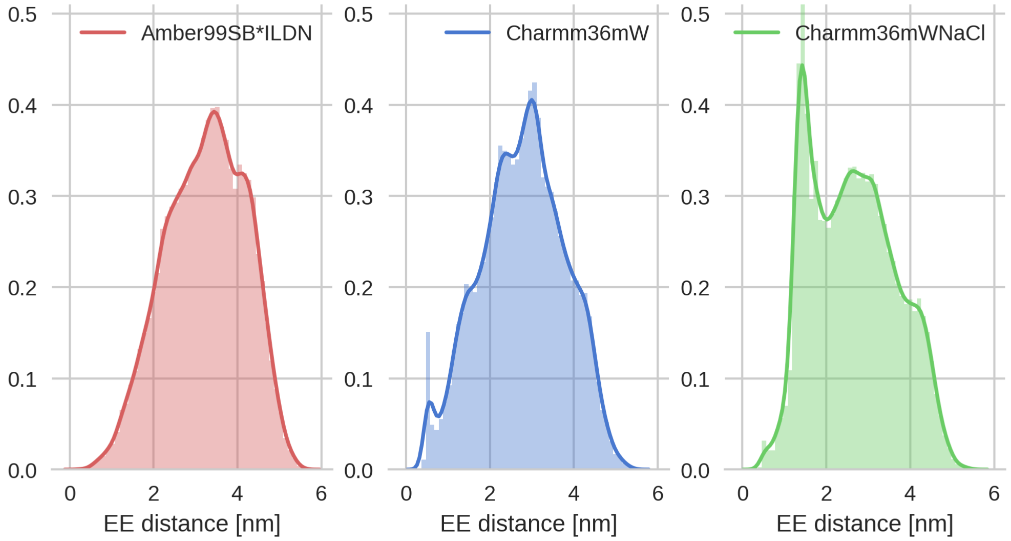


Figure S4: End-to-end distance normalised distributions for the three cases: (A) Amber99SB*-ILDN, (B) Charmm36mW, and (C) Charmm36mW with 150 mM NaCl.

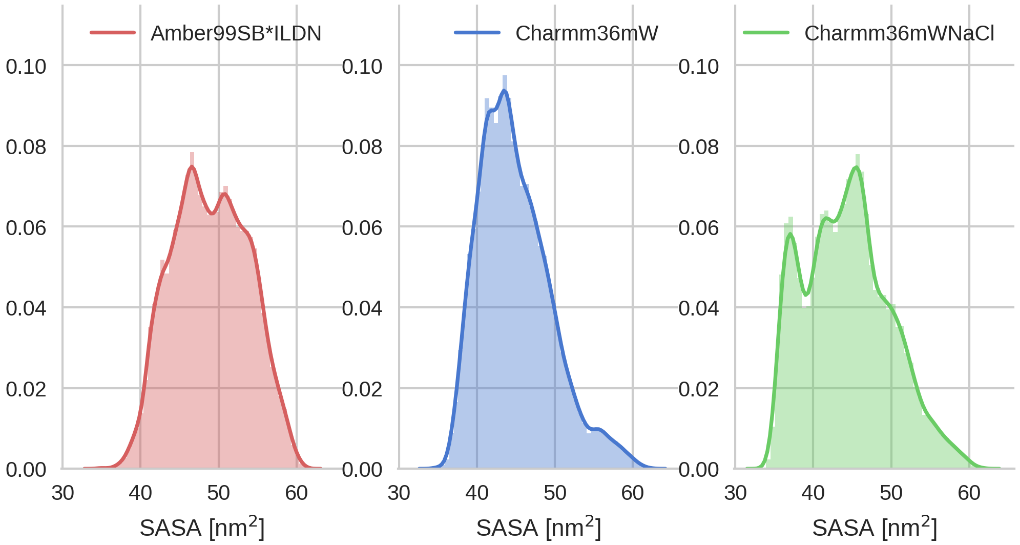


Figure S5: SASA normalised distributions for the three cases: (A) Amber99SB*-ILDN, (B) Charmm36mW, and (C) Charmm36mW with 150 mM NaCl.

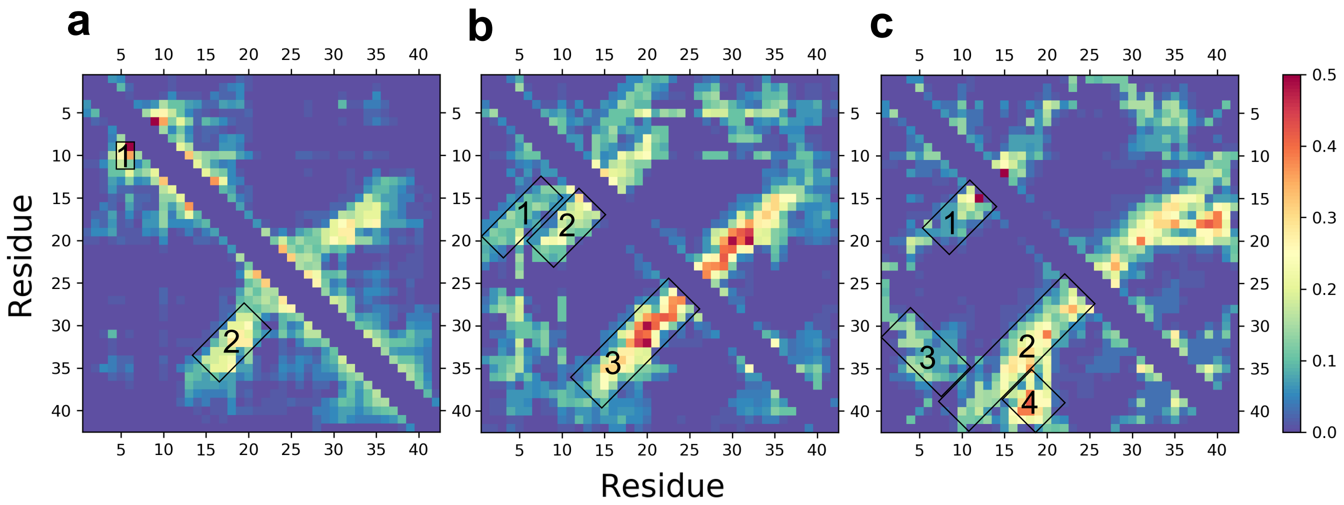


Figure S6: Full contact maps that include the entire analyzed trajectories for the three cases: a) Amber99SB*-ILDN, b) Charmm36mW, and c) Charmm36mW with 150 mM NaCl.

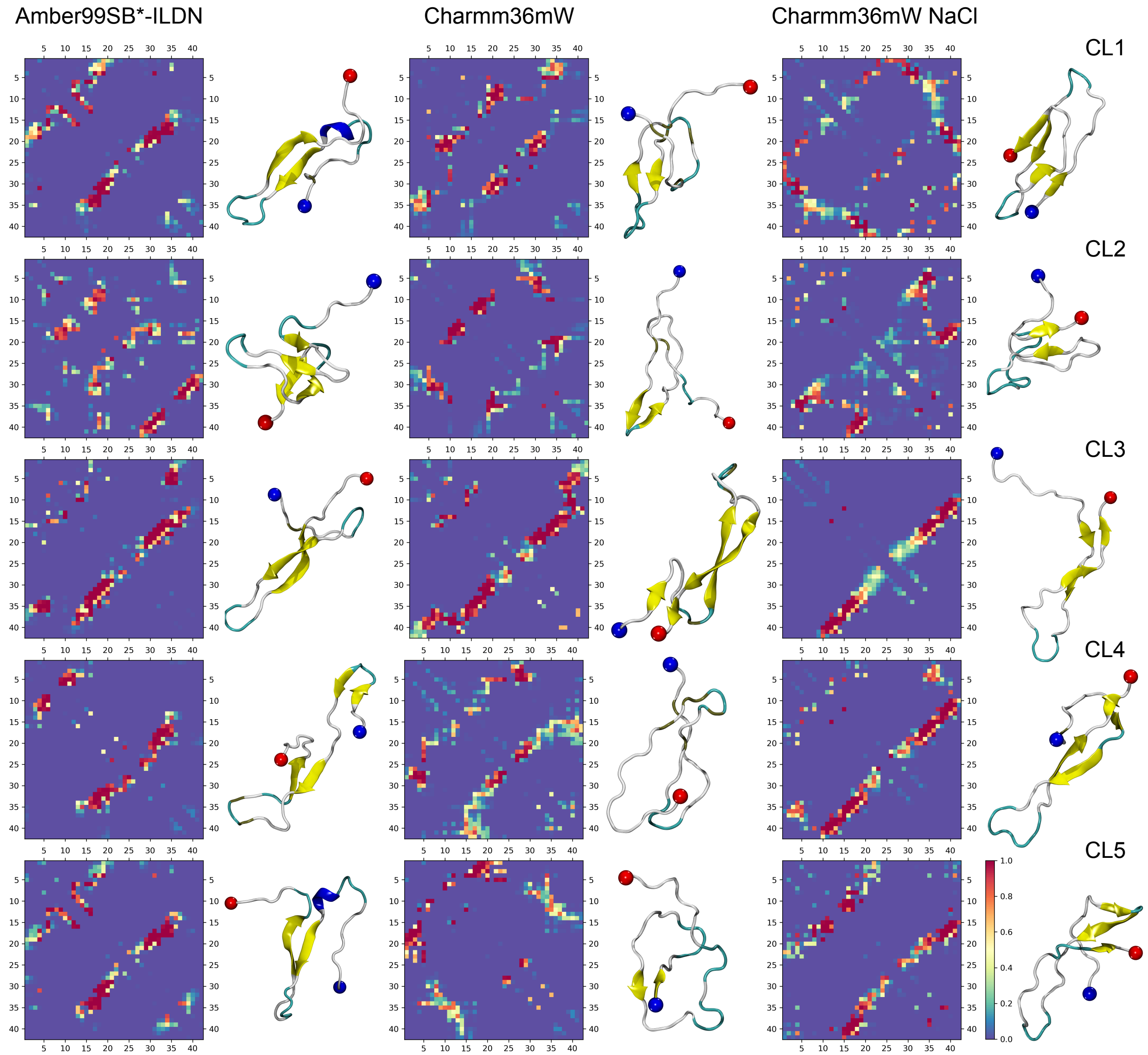


Figure S7: Contact maps and structures for top five clusters with the highest population for the three cases: Amber99SB*-ILDN, Charmm36mW, and Charmm36mW with 150 mM NaCl.

Table ST1: Populations in percentage of the top ten clusters for each simulation with the sum of the populations at the end.

| Cluster | Amber99SB*-ILDN | Charmm36mW | Charmm36mW NaCl |
|---|---|---|---|
| 1 | 3.31 | 7.59 | 14.46 |
| 2 | 2.21 | 7.36 | 7.59 |
| 3 | 2.06 | 5.63 | 2.35 |
| 4 | 1.95 | 5.24 | 2.27 |
| 5 | 1.44 | 1.87 | 1.43 |
| Total | 10.97 | 27.70 | 28.10 |

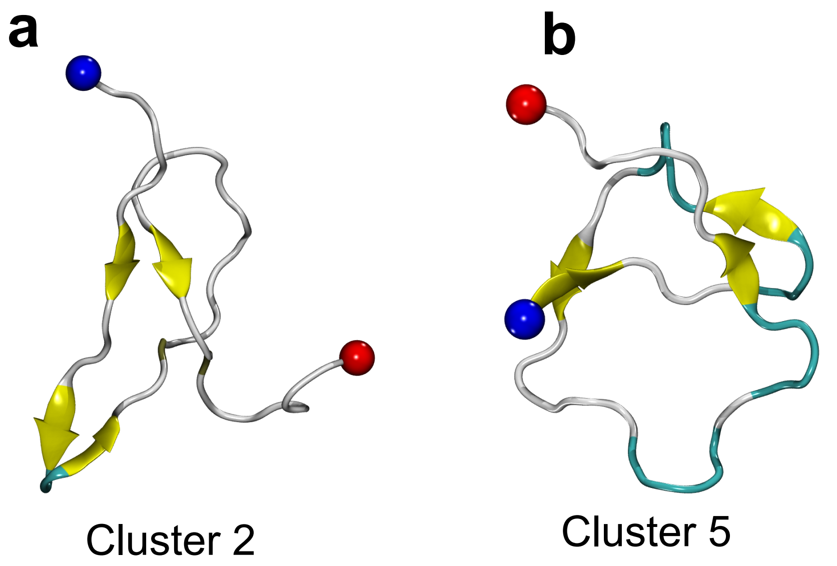


Figure S8: Conformations from clusters two (a) and five (B) from the simulation with Charmm36mW without NaCl that have parallel $\beta$-sheet structure.

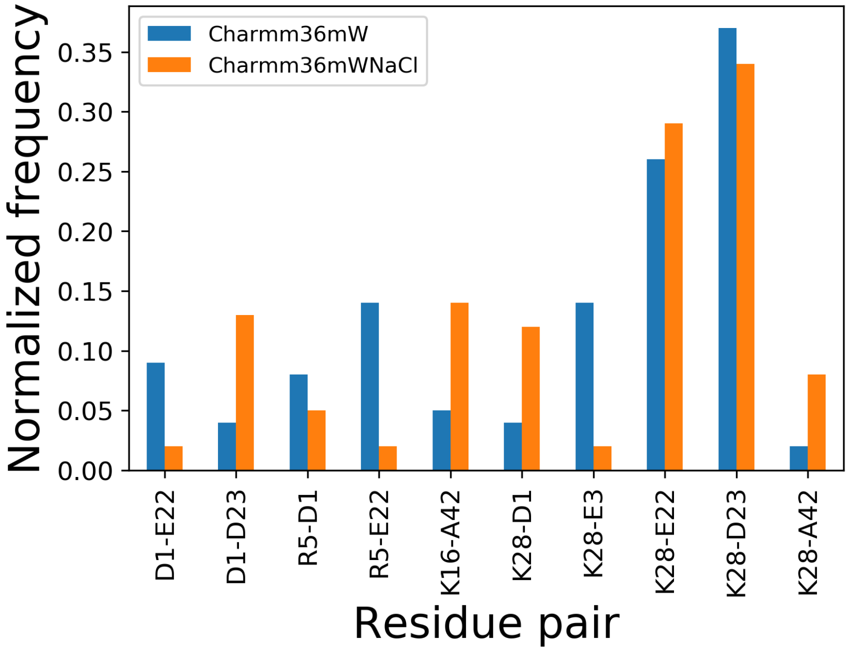


Figure S9: Contact probability between charged amino acids. Only the pairs that have more than 5% contact probability with a cutoff distance of 0.5 nm are shown.

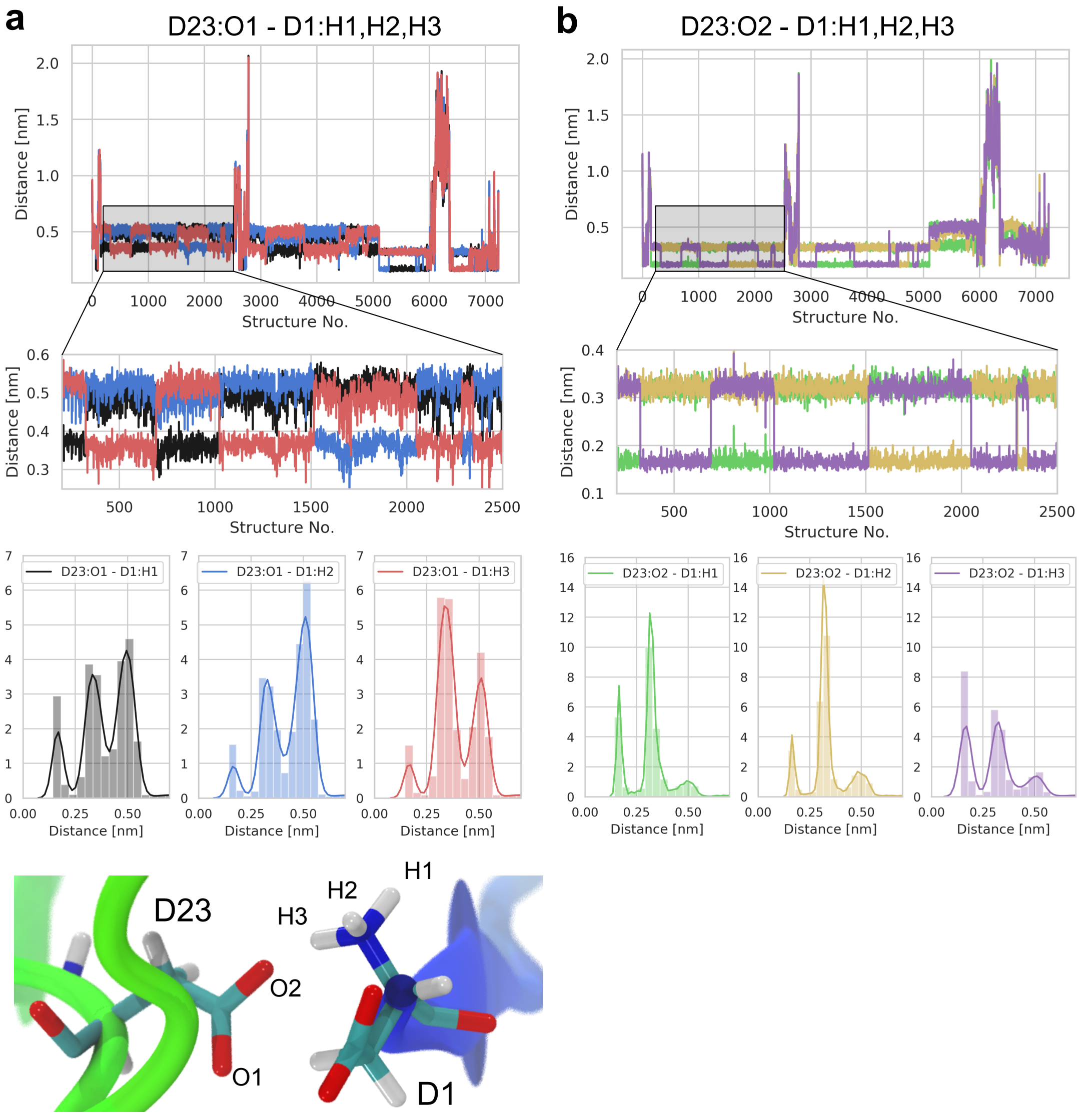


Figure S10: Salt-bridge distances for D23–D1. a) Top - Distance between the oxygen atom O1, see Bottom snapshot, of group COO- of residue D23 and the hydrogen atoms of group NH3+ of residue D1 for all the conformations in cluster one of the simulation with Charmm36mW and NaCl. Middle - Short interval from the plot in a) which highlights the swaping of hydrogens for the shortest distance with the oxygen atom. Bottom - Normalized distributions for distances between the oxygen atom O1 and the three hydrogens. Bottom snapshot - Licorice representation of the amino acids D23 and D1 and the atoms forming the salt-bridge. b) Same as a) but for distances between oxygen O2 of group COO- of residue D23 and the hydrogen atoms of group NH3+ of residue D1.

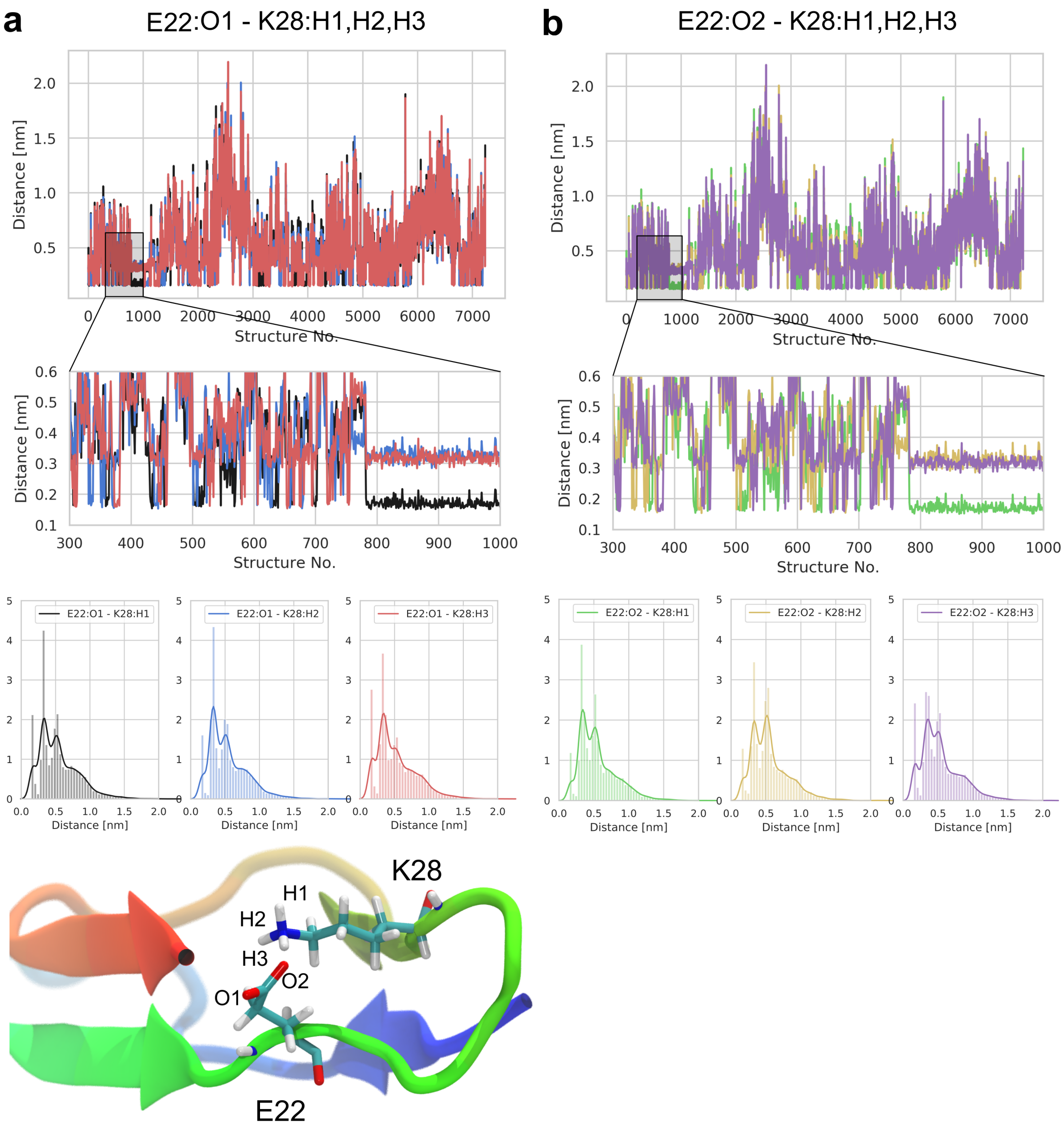


Figure S11: Salt-bridge distances for E22–K28. a) Top - Distance between the oxygen atom O1, see Bottom snapshot, of group COO- of residue E22 and the hydrogen atoms of group NH3+ of residue K28 for all the conformations in cluster one of the simulation with Charmm36mW and NaCl. Middle - Short interval from the plot in a) which highlights the swaping of hydrogens for the shortest distance with the oxygen atom. Bottom - Normalized distributions for distances between the oxygen atom O1 and the three hydrogens. Bottom snapshot - Licorice representation of the amino acids E22 and K28 and the atoms forming the salt-bridge. b) Same as a) but for distances between oxygen O2 of group COO- of residue E22 and the hydrogen atoms of group NH3+ of residue K28.

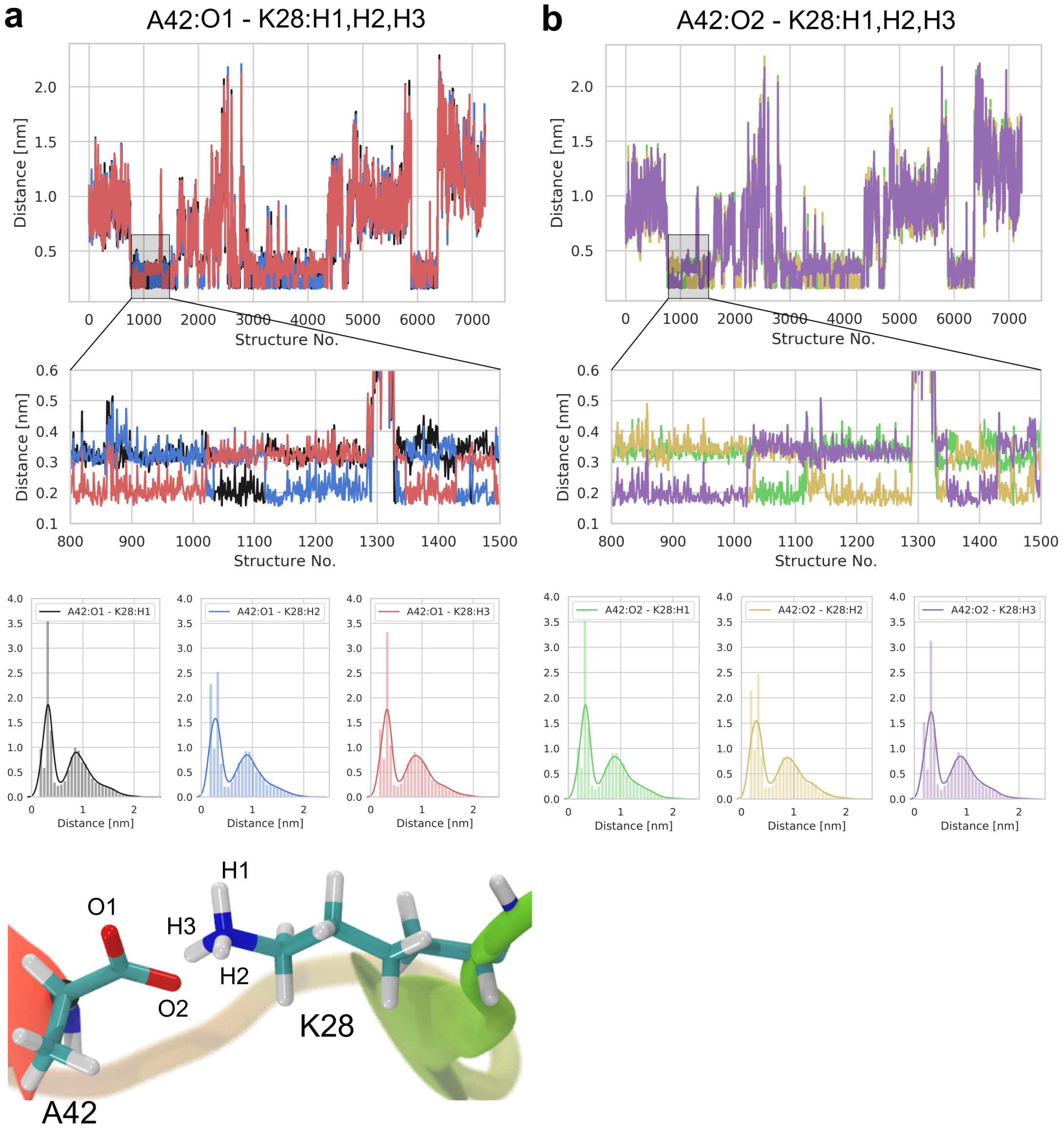


Figure S12: Salt-bridge distances for A42–K28. a) Top - Distance between the oxygen atom O1, see Bottom snapshot, of group COO- of residue A42 and the hydrogen atoms of group NH3+ of residue K28 for all the conformations in cluster one of the simulation with Charmm36mW and NaCl. Middle - Short interval from the plot in a) which highlights the swaping of hydrogens for the shortest distance with the oxygen atom. Bottom - Normalized distributions for distances between the oxygen atom O1 and the three hydrogens. Bottom snapshot - Licorice representation of the amino acids A42 and K28 and the atoms forming the salt-bridge. b) Same as a) but for distances between oxygen O2 of group COO- of residue A42 and the hydrogen atoms of group NH3+ of residue K28.
